# Prebiotically-relevant low polyion multivalency can improve functionality of membraneless compartments

**DOI:** 10.1101/2020.02.23.961920

**Authors:** Fatma Pir Cakmak, Saehyun Choi, McCauley O. Meyer, Philip C. Bevilacqua, Christine D. Keating

**Author notes:** These authors contributed equally to the work (Saehyun Choi, McCauley O. Meyer).

## Abstract

Multivalent polyions can undergo complex coacervation, producing membraneless compartments that accumulate ribozymes and enhance catalysis, and offering a mechanism for functional prebiotic compartmentalization in the origins of life. Here, we evaluated the impact of low, prebiotically-relevant polyion multivalency in coacervate performance as functional compartments. As model polyions, we used positively and negatively charged homopeptides with one to 100 residues, and adenosine mono-, di-, and triphosphate nucleotides. Polycation/polyanion pairs were tested for coacervation, and resulting membraneless compartments were analyzed for salt resistance, ability to provide a distinct internal microenvironment (apparent local pH, RNA partitioning), and effect on RNA structure formation. We find that coacervates formed by phase separation of the relatively shorter polyions more effectively generated distinct pH microenvironments, accumulated RNA, and preserved duplexes. Hence, reduced multivalency polyions are not only viable as functional compartments for prebiotic chemistries, but they can offer advantages over higher molecular weight analogues.

## Introduction

A crucial step in the transition from nonliving prebiotic building blocks to life is the formation of simple protocells from a collection of functional protobiomolecules.^1,2^ The presumed scarcity of functional protobiomolecules, such as ribozymes in an RNA World hypothesis,^3^ poses challenges for passive encapsulation by amphiphile self-assemblies such as vesicles, particularly in the absence of binding interactions to help facilitate encapsulation.^1,4,5^ An alternative physical mechanism for compartmentalization is suggested by the likely presence of nonfunctional oligomeric or polymeric molecules in greater quantities than the protobiomolecules. Associative interactions between these components could drive phase separation, providing membraneless liquid microcompartments to accumulate protobiomolecules, forming a "protocytoplasm".^6^ For example, a type of associative phase separation called complex coacervation occurs readily in solutions of oppositely-charged polyelectrolytes, forming a dense polymer-rich coacervate phase and a dilute continuous phase.^7,8^ The ion pairing interactions between coacervate-forming polymers are nonspecific and hence versatile and achievable with a wide range of biological and nonbiological chemistries.^7,9–12^ Biomolecules such as RNAs can be concentrated within coacervates to orders of magnitude higher than in the external milieu.^9,13^ These higher local concentrations can provide rate enhancements for catalytic RNAs encapsulated within coacervate droplets.^5,14,15^ Coacervates also provide a distinct microenvironment that can differ from the dilute phase in terms of solvent polarity,^16,17^ concentrations of metal ions such as Mg^2+^,^13^ or the presence of cofactors such as spermine, which can enhance ribozyme function.^14,18^ Most studies of coacervation have focused on molecules of high multivalency, to maximize intermolecular interactions.^11,19^ Although complex coacervate formation has been reported from combinations of short cationic peptides (<10 monomers) and nucleotides,^17^ the impact of this reduced multivalency on compartmentalization function has not been explored. The simplicity of lower molecular weight molecules makes them more prebiotically relevant, but their greater translational entropy reduces their propensity to undergo coacervation and impacts phase composition.^11,20,21^ Here, we investigate the functional consequences of using lower multivalency polyelectrolytes to produce membraneless compartments. Suprisingly, we find that membraneless compartments formed using the shorter molecules are functionally superior for some properties important for prebiotic compartmentalization.

## Results

Our model polyelectrolytes were cationic and anionic homopeptides in a range of lengths, as well as anionic mono-, di-, and triphosphates of adenosine (AMP, ADP, and ATP) (Fig. 1A). Although prebiotic peptides would have had greater sequence complexity and length polydispersity, using these simpler peptides allows us to better isolate the effect of multivalency. Charge-matched cationic and anionic components were mixed and the resulting solution classified as containing coacervates, aggregates, or neither (Fig. 1B). A low ionic strength buffer of 15 mM KCl, 0.5 mM MgCl_2_, and 10 mM Tris (pH 8.0) was used to support complex coacervation between low molecular weight peptide or nucleotide components. We then chose polyions that form coacervates over a range of lengths to examine how their multivalency impacts the physical properties of their resulting compartments and how they differ in their ability to both accumulate RNA and influence its base pairing/folding.

**Figure 1.**
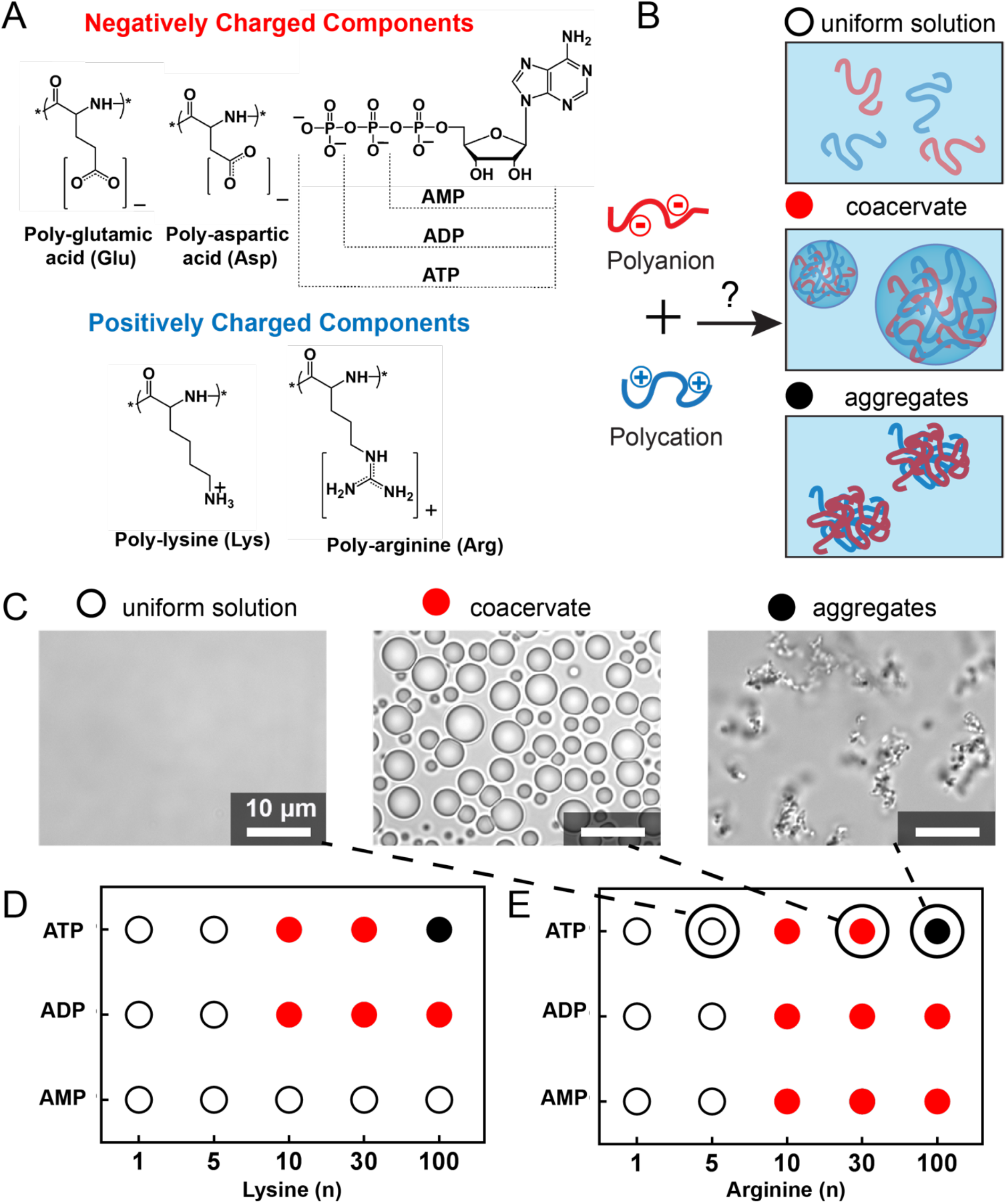
A complex coacervate library was generated by combining pairs of oppositely-charged peptides and nucleotides. (**A**) Structures of negatively and positively charged molecules used as polyanions and polycations in evaluation of coacervation (the nucleotide terminal phosphates are dianionic at pH 8.0 with pKa≈ 1 and ≈ 6. 8). (**B)** Combination of positively and negatively charged components led to a uniformly mixed solution, coacervation, or aggregation, respectively, depending on the details. (**C**) Optical microscope images illustrating samples categorized as uniform solution 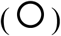, coacervates 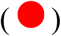 and aggregates 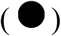; these particular samples are ATP with (Arg)_n_ (n = 5, 30 and 100, left-to-right), corresponding to the points highlighted in panel E. Summary of findings for **(D)** (Lys)_1-100_ and **(E)** (Arg)_1-100_ with AMP, ADP and ATP. The x- and y-axes go in the direction of increasing charge.

### Coacervate formation

We began by determining the shortest length of (Lys)_n_ or (Arg)_n_ (n = 1, 5, 10, 30, or 100) able to form coacervates with each of the nucleotides. Possible outcomes upon mixing the oppositely-charged polyions were uniform solution, coacervate droplets, or solid aggregates (Fig. 1C and Supplementary Fig. 1), with coacervation anticipated for ion pairing interactions able to drive numerous, dynamic intermolecular binding interactions but not so strong as to produce solids.^11,22–24^ The overall pattern in the data is as anticipated, with no phase separation observed for the least-multivalent combinations (lower lefthand corners of Fig. 1D and 1E), coacervates forming at intermediate charge per molecule, and aggregates occurring for the combinations of greatest charge/molecule (i.e ATP with the 100-mer peptides). The shortest peptides that formed coacervates were n = 10 (third columns), which held for both oligolysine and oligoarginine. (Lys)_10_ formed coacervates with ATP and ADP, while (Arg)_10_ also formed coacervates with AMP (Fig. 1D-E, Supplementary Fig. 2). Coacervation with AMP held out even to (Arg)_100_, which can be understood in terms of strong cation-pi interactions possible for Arg-adenosine.^10,25,26^

We then examined coacervate formation between cationic peptides, (Lys)_n_ or (Arg)_n_, and anionic peptides, (Asp)_n_ or (Glu)_n_, as a function of multivalency (Fig. 2). Similar trends were observed for all four combinations, with no phase separation for combinations of the shortest oligomers, and aggregation observed for many combinations of the longest oligomers. Notably, (Arg)_100_ was particularly prone to aggregation, forming solids with even relatively short polycarboxylates (n≥5). The shortest oligomer pairs able to form coacervates under these conditions had at least one component with n=10, and the other with n≥5. For example, (Lys)_10_/(Asp)_5_, (Arg)_10_/(Asp)_5_ or (Arg)_5_/(Asp)_10_, and (Arg)_10_/(Glu)_5_ formed coacervates (Fig. 2). All combinations of (Asp)≥5 and (Lys)≥10 formed coacervates (Fig. 2A). For coacervates containing (Glu)_n_ as the polyanionic component, coacervation occurred at only a few length combinations, and in some cases was accompanied by aggregates that formed in the same samples (Fig. 2C-D). Taken together, the data presented in Fig. 1 and 2 demonstrate that rather short polyelectrolytes (n=10 polycations plus mononucleotides, or n=5/n=10 polycation plus polyanion combinations) readily form complex coacervates, and provide us with a small library of coacervate compositions across which we can compare compartmentalization.

**Figure 2.**
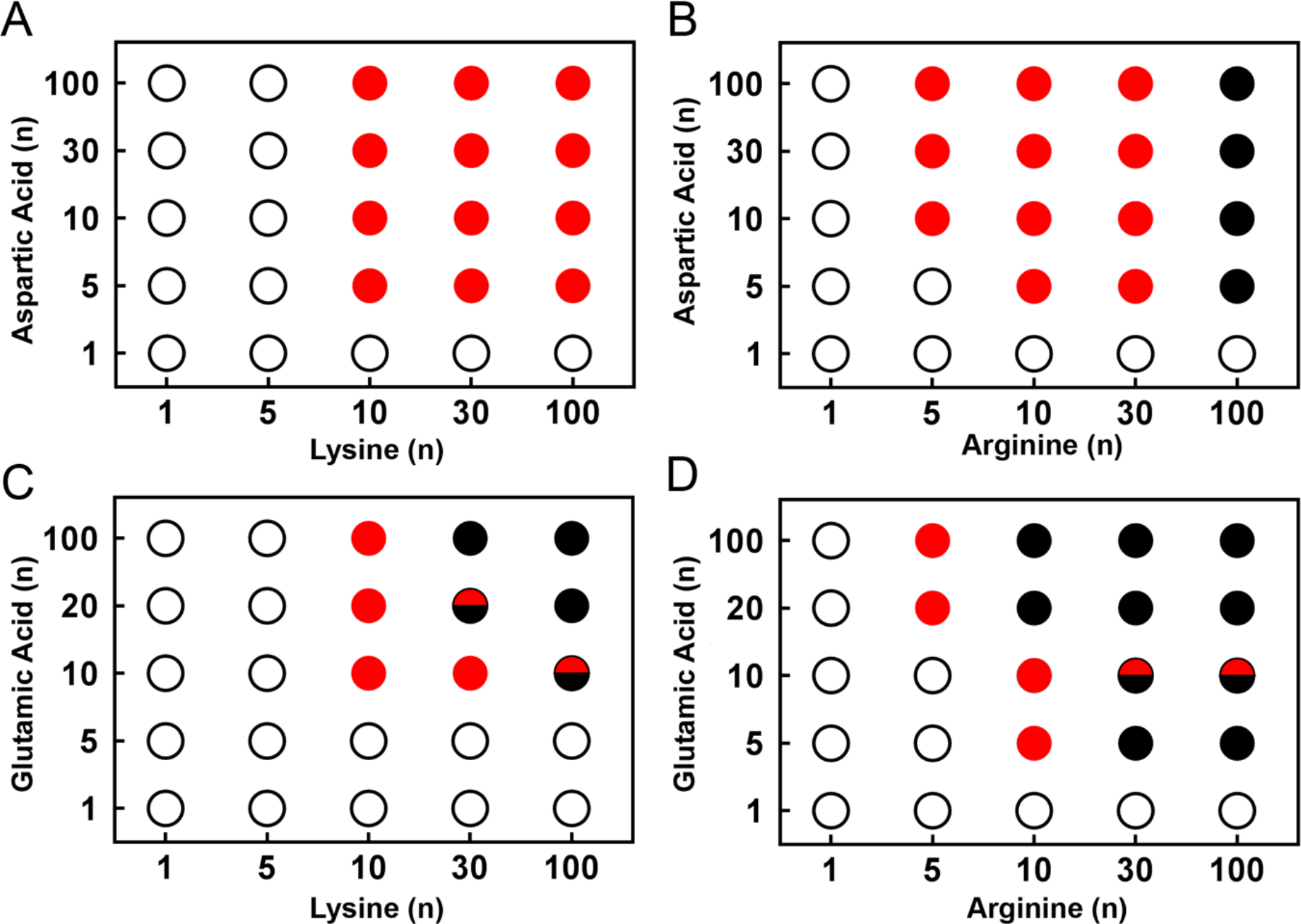
Summary of coacervation results generated by combining pairs of oppositely-charged peptides. Polyanions ((Asp)_n_ or (Glu)_n_) were mixed with polycations ((Lys)_n_ or (Arg)_n_) at 10 mM charge 1:1 charge ratio. Formation result of pairs of peptides of **(A)** (Lys)_n_ / (Asp)_n_ **(B)** (Arg)_n_ /(Asp)_n_, **(C)** (Lys)_n_ / (Glu)_n_, and **(D)** (Arg)_n_ / (Glu)_n_ where n=1, 5, 10, 20, 30 or 100. Symbols indicate observation of uniform solution 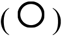, coacervates 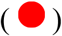, aggregates 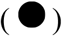 or 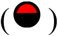 the presence of both aggregates and coacervates together in the same sample.

### Salt dependence of coacervate formation

Ion pairing-based phase separation is strongly dependent on solution ionic strength, with critical salt concentrations above which coacervates dissolve dependent upon multivalency.^19,22,27^ We evaluated the salt stability of coacervates formed using primarily n=10 cationic peptides with anions that included nucleotides and both carboxylate oligopeptides (Fig. 3A-D, Supplementary Table 1). For coacervates formed between nucleotides and both n=10 polycations, salt resistance increased with increasing nucleotide charge found in going AMP to ATP (Fig. 3A–B). Additionally, the (Arg)_10_/nucleotide coacervates had greater salt stability than their (Lys)_10_ counterparts, >600 mM KCl for (Arg)_10_/ATP but only ~ 100 mM for (Lys)_10_/ATP. Similar trends were apparent in coacervates formed with combinations of cationic and anionic peptides of various lengths, although they are less pronounced (Fig. 3C-D). Coacervates with longer peptides had higher salt resistance as expected for greater multivalency.^11,24,27^ We again observe that coacervates formed with oligoarginines have markedly higher salt stability than those formed with oligolysines; for example, (Arg)_10_/(Asp)_10_ is stable to nearly 1.5 M KCl, while (Lys)_10_/(Asp)_10_ coacervates dissolve above ~ 300 mM KCl (Supplementary Table 1). In comparing anionic oligopeptides, we see greater salt stability for oligoaspartates than oligoglutamates, when all else is equal (Fig. 3C-D). Molecule-specific differences in salt stability reflect differences in interactions of polyions with each other, themselves, and/or solvent. For example, the possibility of cation-pi interactions, as well as differences in hydrogen bonding and/or polymer hydrophobicity can influence salt stability. We attribute the surprisingly high salt stability of (Arg)_10_/nucleotide coacervates to cation-pi binding, which is known to be strong for Arg residues and adenosine nucleobases.^25,26,28^

**Figure 3.**
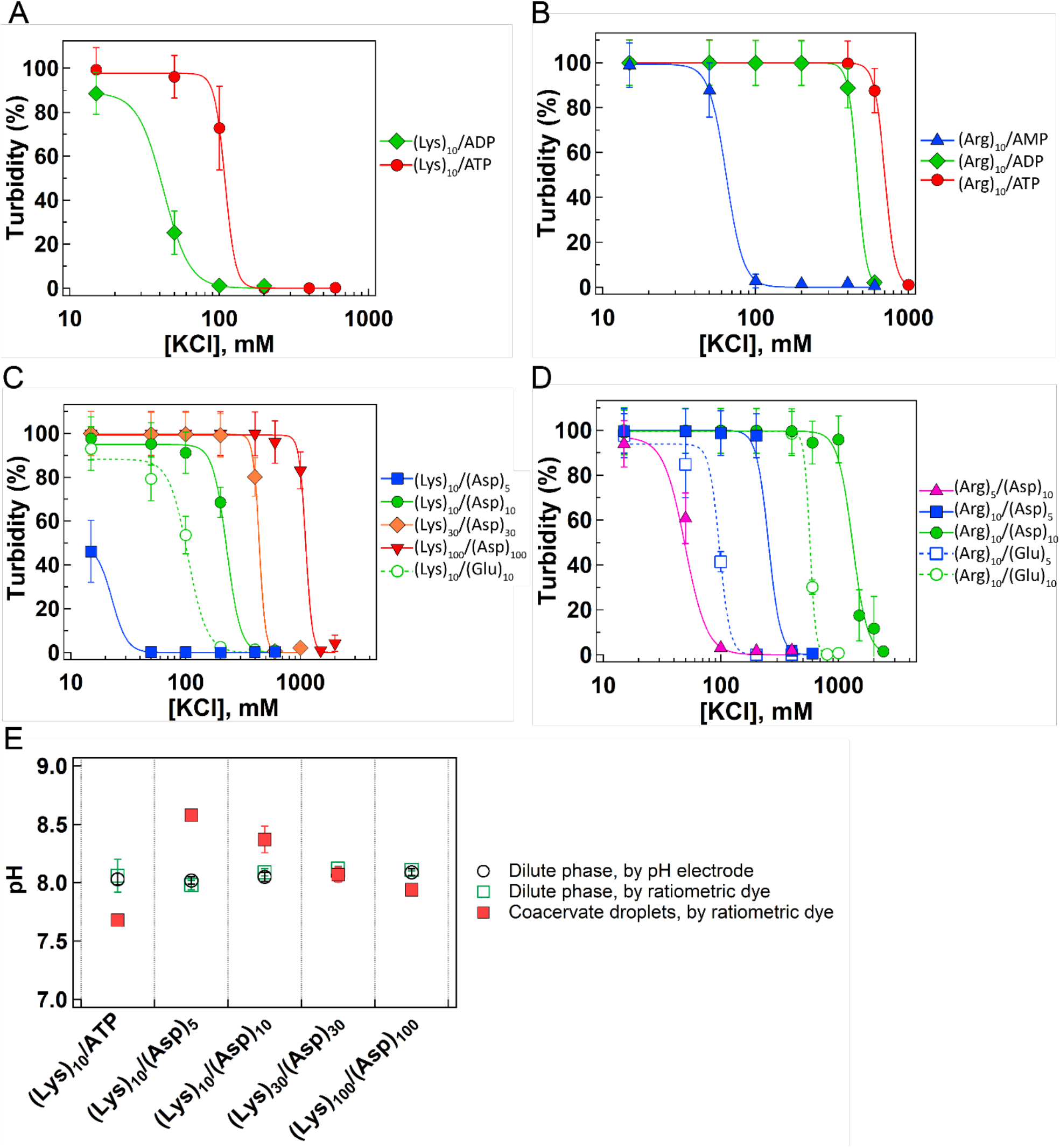
Physical properties for selected coacervate systems as nucleotide charge and peptide length increase. **(A)** Salt resistance of coacervates formed from (Lys)_10_ and nucleotides (ADP and ATP). **(B)** Salt resistance of coacervates formed from (Arg)_10_ and nucleotides (AMP, ADP and ATP). **(C)** Salt resistance of coacervates formed from Lys_10-100_ as length of (Asp)_n_ increases (n=5, 10, 30 and 100), and with Glu_10_. **(D)** Salt resistance of coacervates formed from Arg_10_ and Asp_5,10_ or Glu_5,10_. Critical salt concentrations determined from the fits for curves in panels A-D are available in Supplementary Table 1. Error bars show standard deviation of measurements over at least three independent samples. Relative errors are minimally 10% turbidity values in panels A-D, and may not be visible on low turbidity values. **(E)** Measured (black circle) and calculated (green box) pH of the dilute phase, and calculated pH of coacervate droplets (red box) for different coacervate systems. See Supplementary Fig. 3 and 4 for calculations.

Early Earth conditions are thought to have encompassed a wide range of salt concentrations, from ponds to ocean water.^29,30^ Our results indicate that even at just n=10, certain oligopeptide-based coacervates persist even above 1 M ionic strength, supporting the relevance of coacervate-based prebiotic compartments to diverse prebiotic scenarios extending beyond freshwater to brackish waters, oceans or submarine hydrothermal vent systems alike.

### pH inside coacervates

Coacervates contain high concentrations of their component molecules, with cationic and anionic functional groups present up to molar levels.^13,17,19^ We reasoned that high local concentrations of amine, carboxylate, and/or phosphate moieties could impact proton availability inside coacervates. We evaluated this possibility using the pH sensitive ratiometric dye C-SNARF-1 to measure apparent pH inside coacervate droplets. In the case of pH for the dilute phase, we also directly measured the pH with an electrode, where we obtained very similar values to those measured with the dye, validating in-coacervate measurements (Fig. 3E, compare black circles and green squares). Coacervates formed from (Lys)_10_/ATP and (Lys)_n_/(Asp)_n_ peptide pairs were chosen for pH measurements owing to their ability to form coacervates over a wide range of oligopeptide lengths (Supplementary Fig. 2). We observed differences in apparent pH between the coacervate droplets and the external continuous phase for several of the systems (Fig. 3E). The pH of (Lys)_10_/ATP coacervates was ~7.7 (more acidic than dilute phase), likely reflecting the high local ATP concentration (*γ* phosphate pKa ≈ 6.8). (Lys)_10_/(Asp)_5_ coacervates had an apparent local pH near pH 8.5, ~0.5 pH units *higher* than the dilute continuous phase, which we interpret as resulting from excess of amine moieties within the coacervate owing to the greater multivalency of (Lys)_10_ as compared to (Asp)_5_. The other systems tested here had more similar apparent local pH inside and outside the coacervate droplets, with internal apparent pH decreasing as multivalency increased from n=10 to 100; the cause of this apparent trend is unclear but could be related to increases in local concentration of both polymers as multivalency is increased, and/or to the presence of proportionately less N- and C-terminal moieties. Overall these data demonstrate that self-assembly of even relatively primitive, simple polyions can provide compartments with local pH that differs from the external milieu; indeed, the largest pH differences were observed for the coacervates formed from the smallest polyions.

### RNA partitioning in coacervate systems

To serve as functional prebiotic compartments, coacervate droplets should concentrate solutes of interest. We therefore sought to determine the impact of reduced polyion multivalency on the ability of coacervates to accumulate RNA oligonucleotides. Fluorescently-labeled RNA designed to exhibit minimal secondary structure was used for partitioning studies (see Methods). When fluorescently labeled 10 nt ssRNA was added at a final concentration of 0.1 µM to coacervate containing systems, significantly higher fluorescence was observed inside the coacervate droplets as compared to the continuous phase for all systems except (Lys)_100_/(Asp)_100_ (Fig. 4A-B and Supplementary Table 2). We saw little difference between the RNA concentration within droplets formed from (Lys)_10_/(Asp)_5_, (Lys)_10_/(Asp)_10_ and (Lys)_30_/(Asp)_30_, with each of these systems showing at least 100-fold increase in local RNA concentration, to ~ 11 µM inside the droplets (K_RNA 10mer_ = 160 - 400). The (Lys)_10_/ ATP coacervates had even higher local concentration of 10mer ssRNA (~ 43 µM, K_RNA 10mer_ = 5.3 x10^3^) (Fig. 4C).

**Figure 4.**
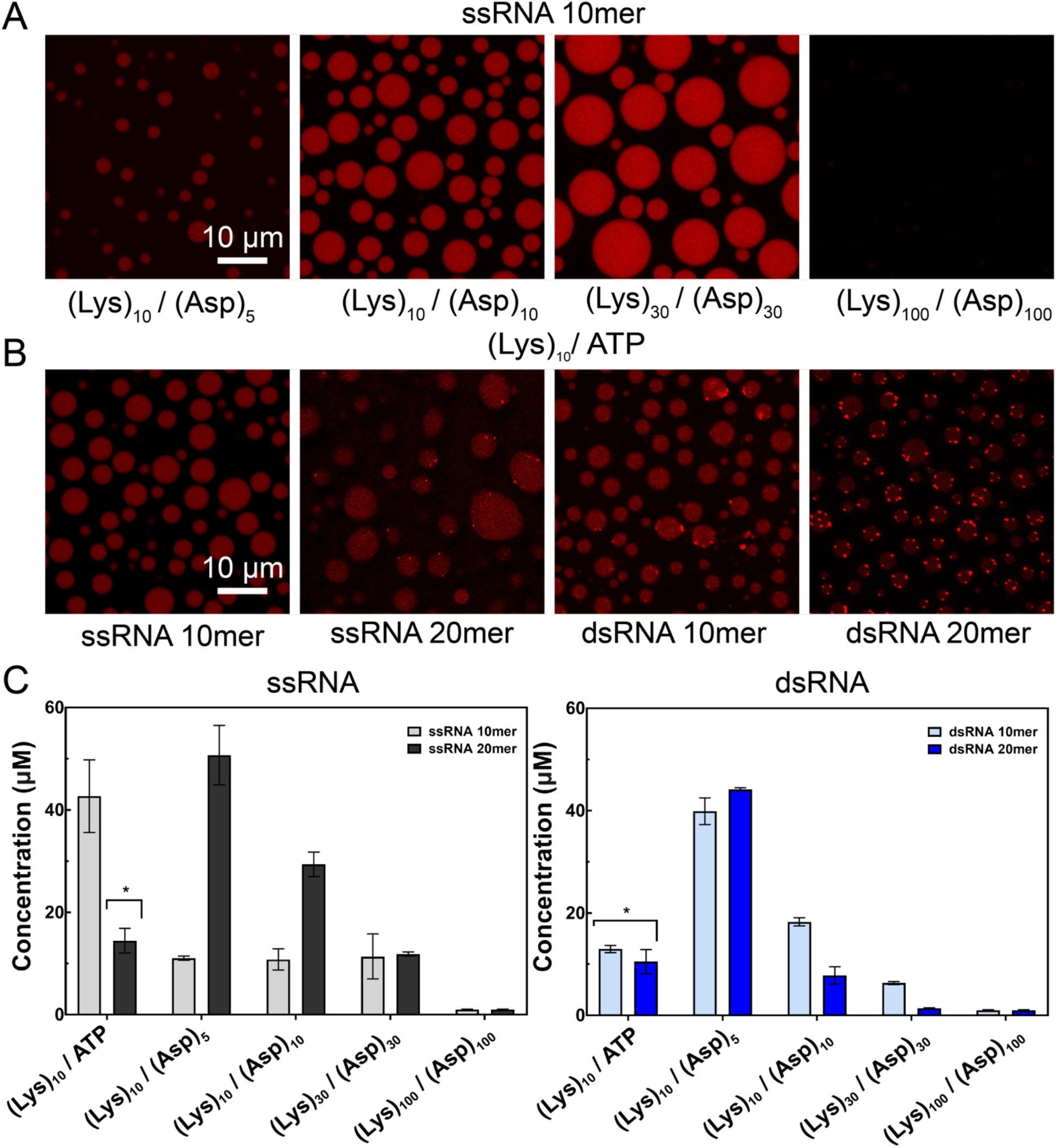
Partitioning of ss and dsRNA 10- and 20-mers in selected coacervates. **(A)** Fluorescence images of Cy3-labeled ssRNA 10mer in (Lys)_10_/(Asp)_5_, (Lys)_10_/(Asp)_10_, (Lys)_30_/(Asp)_30_ and (Lys)_100_/(Asp)_100_ coacervate systems. **(B)** Fluorescence images of Cy3 labeled ssRNA 10mer, ssRNA 20mer, dsRNA 10mer and dsRNA 20mer all in a (Lys)_10_/ATP coacervate. Note that laser intensity was optimized separately for each sample in panels A and B; quantification based on calibration curves is shown in panel C. **(C)** Calculated concentration of ssRNA 10mer, ssRNA 20mer, dsRNA 10mer and dsRNA 20mer in the coacervate droplets for (Lys)_10_/ATP, (Lys)_10_/(Asp)_5_, (Lys)_10_/(Asp)_10_, (Lys)_30_/(Asp)_30_ and (Lys)_100_/(Asp)_100_ coacervate pairs. Error bars show the standard deviation of multiple individual droplets obtained over at least three independent samples. Note that aggregation was observed for (Lys)_10_/ATP for partitioning of ssRNA 20mer, ds RNA 10mer and dsRNA 20mer. “*” represents the coacervates containing speckles. RNA strands were added to a final concentration of 0.1 µM for all cases. RNA concentration in the droplets was calculated without including ATP/(Lys)_10_ bright speckles, which we interpret as a new phase in which RNA is the main polyanionic component.

Increasing the ssRNA length from 10 to 20 nucleotides led to significantly higher RNA concentrations in the droplet phase for (Lys)_10_/(Asp)_5_ and (Lys)_10_/(Asp)_10_, with ~ 50 µM in (Lys)_10_/(Asp)_5_ coacervates, a 500-fold increase over the average RNA concentration in the total sample (Fig. 4 C and Supplementary Table 2). However, as the length of the coacervate components increased, the concentration of the ssRNA 20mer in the droplet phase decreased markedly, dropping to ~30 µM and then ~12 µM for (Lys)_10_/(Asp)_10_ and (Lys)_30_/(Asp)_30_, respectively. As with the 10mer, there was no preferential partitioning of the 20mer into (Lys)_100_/(Asp)_100_ droplets. The partitioning trends are consistent with a competitive displacement RNA accumulation mechanism, whereby the RNA enters the coacervate by displacing (Asp)_n_ to interact with (Lys)_n_. As the multivalency of the oligoaspartate increases, it becomes more difficult for the RNA to compete with it for binding sites on the oligolysine.^13,15^ By increasing the charge ratio of cationic to anionic groups, RNA accumulation can be encouraged.^15,31^ Indeed, for (Lys)_100_/(Asp)_100_ with a slight excess of (Lys)_100_ (1.2: 1 charge ratio), we achieved ~11 and ~18 µM partitioning for ssRNA 10mer and 20mer, respectively (Supplementary Fig. 5). In (Lys)_10_/ATP samples, speckles of brighter fluorescence intensity were observed within the coacervate droplets for all RNA except ssRNA 10mer (Fig. 4B); these bright puncta are consistent with formation of a second coacervate phase in which RNA is the predominant anionic component.^21,31^ Together, the results shown in Fig. 4 indicate that membraneless compartments formed by phase separation of relatively simple polyions ((Lys)_10_/ATP, (Lys)_10_/(Asp)_5_) provide strong accumulation of ssRNA oligonucleotides. More generally, these data suggest that accumulation of polyions such as nucleic acids in complex coacervates is actually favored by producing these membraneless compartments using shorter polyelectrolytes.

Nucleic acid function often requires base-pairing interactions, and certain membraneless organelle models based on intrinsically disordered proteins have shown preferential partitioning of single-stranded versus double-stranded nucleic acids, particularly for 20-nt and higher lengths, which was ascribed to the greater persistence length of the double-stranded nucleic acids resisting partitioning.^32^ Although the chemistry of our coacervates differs from the Ddx4-based systems, we also observe some length- and strandedness-related differences in RNA accumulation across the different coacervates (Fig. 4C, Supplementary Fig. 6-9, and Supplementary Tables 2 and 3). This discrimination is most notable for coacervate systems with the shortest peptide pairs (Lys)_10_/(Asp)_5_ and (Lys)_10_/(Asp)_10_. For 10-mer RNA, we found that dsRNA showed generally stronger partitioning into the coacervates than ssRNA (Fig. 4C), consistent with greater charge density of dsRNA but differing from how these molecules partitioned in Ddx4 protein-based droplets, where single-stranded nucleic acids had stronger accumulation.^32^ In contrast, the 20-nt RNA showed greater local accumulation of the ss than dsRNA, which may be due to the longer persistence length of the dsRNA 20-mers, which resists partitioning. The concentration of dsRNA 10mer and 20mer in the droplets decreased with increasing length of the (Lys)_n_/(Asp)_n_ coacervate-forming peptide pairs, and no RNA accumulation was seen for the (Lys)_100_/(Asp)_100_ coacervates (Fig. 4). This is much like that observed for ssRNAs, except there only the 20mer showed the coacervate-length dependence (Supplementary Table 2 and 3). RNA-rich speckles of bright fluorescence were observed for all dsRNA experiments, especially for the 20mers, and the (Lys)_10_/ATP coacervate system (Figure 4 B). This could be related to differences in how single- and double-stranded oligonucleotides interact with the polycation, which have been reported in studies of (Lys)_n_/DNA coacervation.^33^

### Effect of coacervate microenvironment on nucleic acid hybridization

Differences in preferential accumulation of ss-versus dsRNAs observed in Fig. 4 suggest that the equilibrium between these two states may differ in coacervates as compared to dilute buffer.^32^ We evaluated the impact of various coacervate microenvironments on nucleic acid hybridization by fluorescence resonance energy transfer (FRET). A 3’-Cy3-labeled RNA 10mer sense strand was allowed to hybridize to a 5’-Cy5-labeled antisense strand to produce dsRNA labeled with the Cy3/Cy5 FRET pair (Supplementary Tables 4-6); loss of FRET signal indicates reduced fraction of dsRNA. FRET was assayed in (Lys)_10_/ATP, (Lys)_10_/(Asp)_10_, (Lys)_30_/(Asp)_30_ and (Lys)_100_/(Asp)_100_ coacervate-forming peptide pairs (Fig. 5A). We observed high FRET signal for RNA inside (Lys)_10_/ATP and (Lys)_10_/(Asp)_10_ coacervates, similar to buffer alone, supporting intact duplex. Decreased FRET efficiency was observed for RNA in coacervates formed from the longer peptides, dropping its maximum value by approximately on-half for (Lys)_30_/(Asp)_30_ and one-third for (Lys)_100_/(Asp)_100_ coacervates. Thus, whether coacervate-encapsulated RNA duplexes are disrupted or retained depends on the oligopeptide multivalency, with helicase-like activity, similar to that reported for Ddx4 coacervates, observed only for n≥30.^32^ In principle, some destabilization of RNA base-pairing in prebiotic compartments could prove useful by allowing misfolded RNAs to refold into functional forms, which could become important as encapsulated RNA lengths increase.

**Figure 5.**
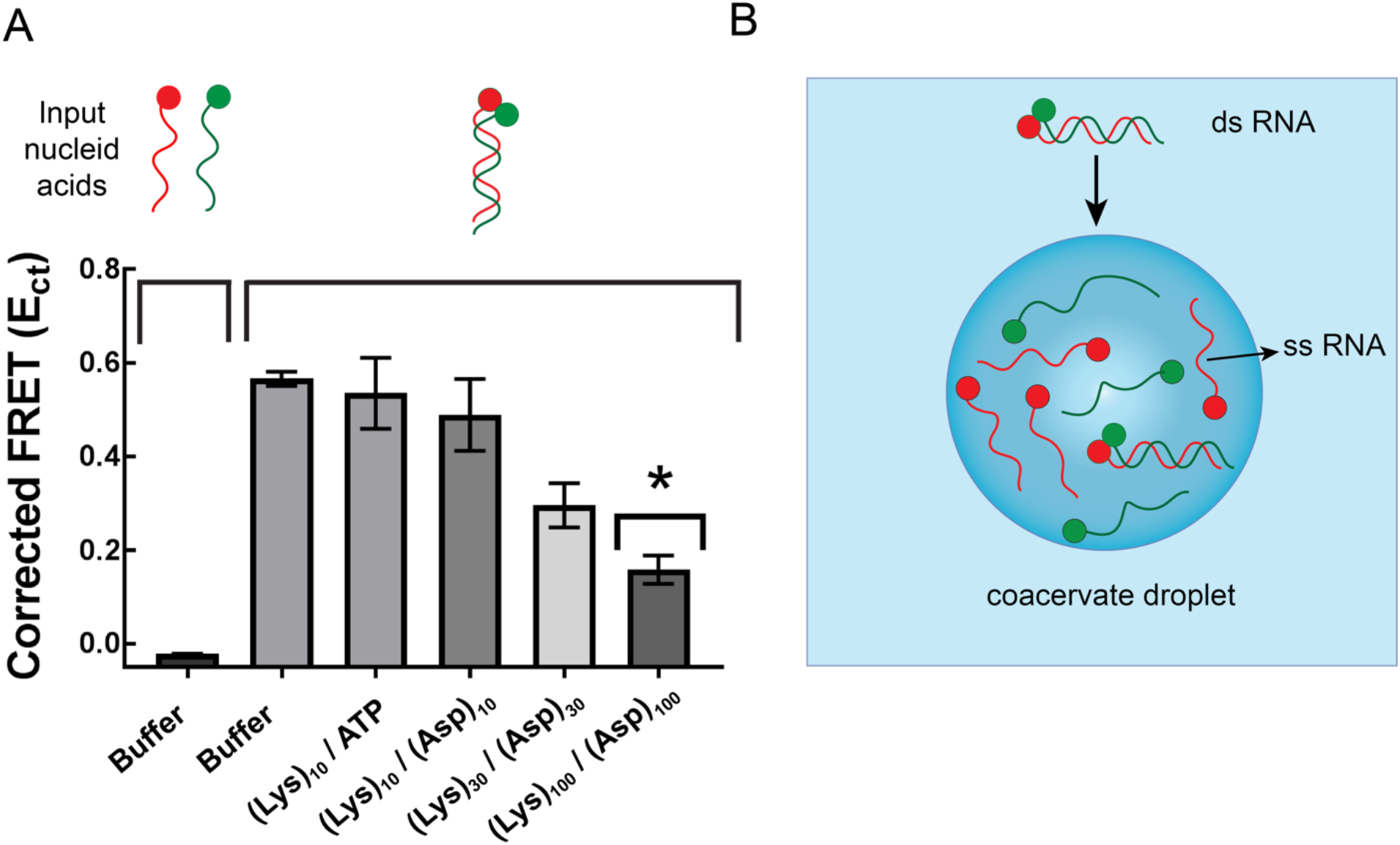
Double helix structure of 10-mer RNA in coacervate droplets. **(A)** Corrected FRET values calculated for buffer and different type of coacervates and buffer. The-FRET single-stranded, noncomplementary control was performed by mixing Cy3 and Cy5 labeled oligonucleotides of the same sequence (see Supplementary Table 5). *Due to low accumulation in the (Lys)_100_/(Asp)_100_ coacervates, a slight excess of the polycation was added (charge ratio 1.2: 1) to increase RNA concentration in the droplets, and the amount of ds RNA added was increased from 0.1 µM to 1 µM. **(B)** Schematic representation of ds RNA 10mer in coacervate systems, which are composed of (Lys)_30_/(Asp)_30_ and (Lys)_100_/(Asp)_100_. Error bars show standard deviation of multiple individual droplets obtained over at least three independent samples.

### Effect of coacervate microenvironment on functional RNA structure

RNAs rely on complex secondary and tertiary structures for function.^34^ We evaluated the impact of compartmentalization inside coacervates on the folding of a T7 transcript, tRNA^phe^, which has extensive secondary structure and a compact and tertiary structure. In-line probing (ILP),^35^ which measures the flexibility of each nucleotide by its ability to undergo hydrolysis, was used to compare the structure of tRNA^phe^ in different coacervates to its native fold in buffer (Fig. 6 and Supplementary Fig. 10; see also ILP control discussion in SI). We found similar reactivity for tRNA in each of the coacervates tested, with some regions remaining folded and others melting out (Supplementary Fig. 11-13). The D-stem, AC-stem and TC-stems remain largely unreactive, supporting retention of the base of the cloverleaf secondary structure. However, the acceptor stem is largely melted out, which is especially clear at the 5’ portion of the acceptor stem (Supplementary Fig. 13). The increase in acceptor stem reactivity could be due to increased breathing of the stem from interaction with the polycations in the coacervate, which is harder to form owing to its long-distance nature.

**Figure 6.**
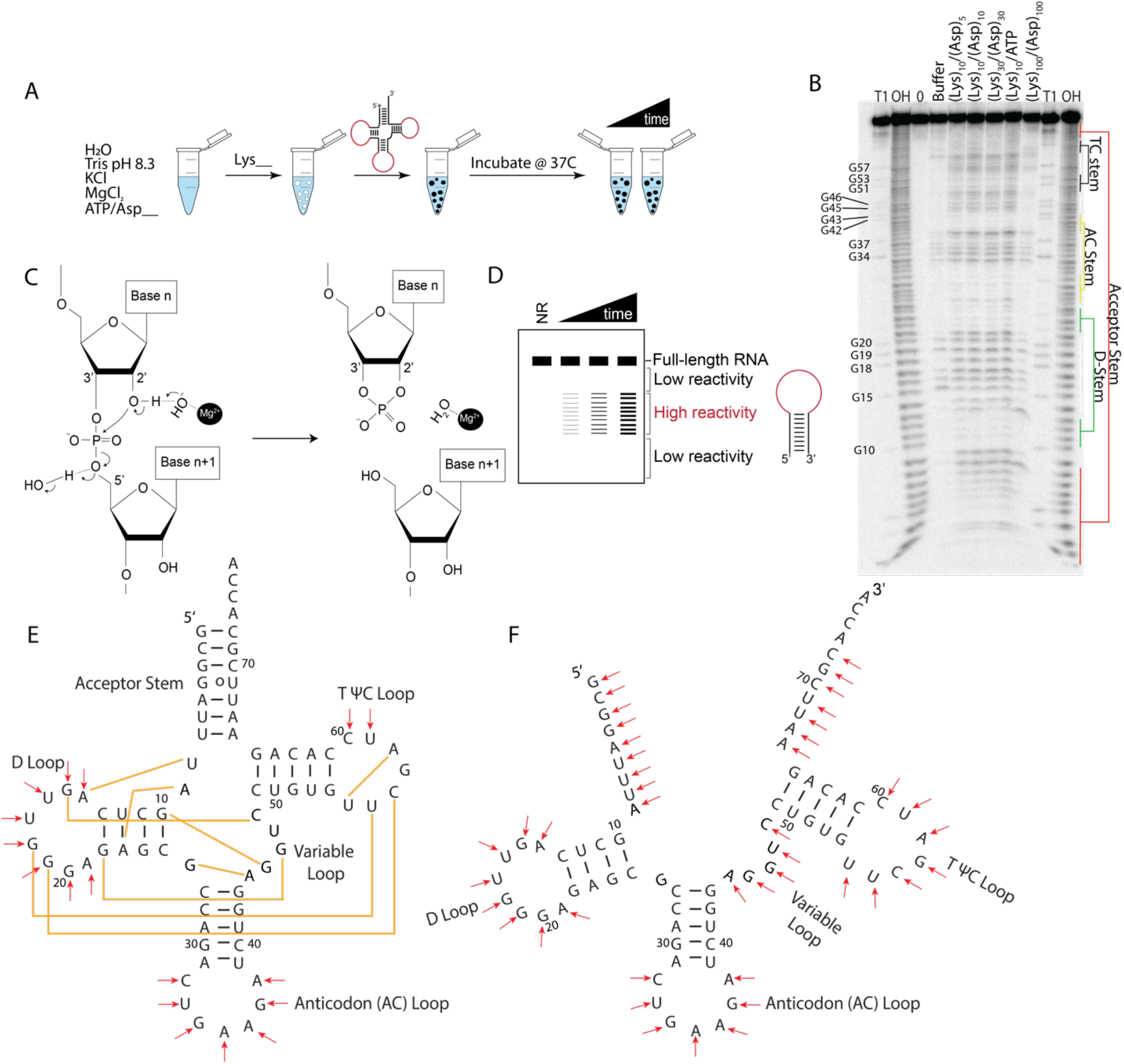
Analysis of RNA structure in coacervates by in-line probing (ILP). (**A**) ILP sample pipeline. (**B**) ILP of tRNA^phe^ in all buffer and coacervate conditions tested. (**C**) Mechanism of ILP strand cleavage of RNA. (**D**) ILP reactivities of radiolabeled RNAs can be read out on denaturing PAGE. (**E**) Natively-folded tRNA^phe^ in the absence of coacervates used as a control to monitor folding within coacervates; yellow lines depict tertiary interactions. (**F**) Structure of tRNA^phe^ in coacervates suggested by data in Panel B, which show that the RNA was unfolded at the acceptor stem in all coacervate conditions evaluated.

We next checked on tertiary structure. Reactivity in the TψC-loop was enhanced in the coacervates, suggestive of some unfolding of tertiary structure (Fig. 6B). Moreover, the D-loop, which is the tertiary structure partner of the TψC loop, also gains in reactivity, especially at G18, which has a known interaction with U55 (Fig. 6E). Further supporting unfolding of the tertiary structure, reactivity is increased in the variable loop. Taken together, these data indicate that short range secondary structure was maintained, while long-range secondary structure was denatured; moreover, tertiary structure was denatured in the presence of all coacervates. (Fig. 6 F). Although ILP for the tRNA^phe^ revealed little difference between the coacervates, FRET studies for the 10-mer duplex indicated denaturation only in higher-multivalency coacervates (Fig 5A). We note that while tRNA^phe^ tertiary structure was not well populated in these coacervates, we have demonstrated ribozyme activity, which is dependent on tertiary structure, in related coacervates.^5,14,15^ One possibility is that RNA tertiary structure forms in a minor population of RNAs that is responsible for activity.

## Discussion

Associative phase separation is an organizational mechanism in extant cells, leading to membraneless organelles enriched in proteins and nucleic acids.^36,37^ The spontaneous occurrence of coacervates in diverse macromolecular systems, and their ability to accumulate solutes such as RNA, suggests possible roles in prebiotic compartmentalization and protocell formation.^2,6^ Since lower molecular weight,^38^ and hence lower multivalency, oligomers are likely more relevant to prebiotic scenarios than larger macromolecules, we considered whether such coacervates could offer sufficiently distinct microenvironments to enable RNA accumulation and function. We find that coacervates formed by low-multivalency components can match or exceed performance of those formed by higher-multivalency counterparts in all areas tested here except salt resistance.^27^ Notably, the lower-multivalency coacervates were more effective at maintaining distinct pH and accumulating RNA. This has profound implications for early Earth, when functional protobiomolecules would have been scarce amongst a complex milieu of molecular components, many of which may have had similar chemistries. In such a scenario, the compartments formed by these relatively low-multivalency polyions could have accumulated longer, potentially functional protobiomolecules (e.g. RNAs) to relatively high local concentrations.

The changes in RNA structure inside complex coacervates observed here, which were polyion length-dependent for the FRET RNA 10-mer duplex formation studies but not for the ILP tRNA^phe^ studies, warrant further investigation and point to the likelihood of coacervate- and RNA-specific impacts on RNA folding in these membraneless compartments. Such findings are also of interest in light of the central role of RNA in the membraneless organelles of extant biology.^36^

## Materials and Methods

### Coacervate preparation

Coacervate samples were prepared in charge concentration ratios of 10 mM (charge of the molecule x its concentration = charge concentration) with a total volume of 100 μL in HPLC grade water and 10 mM Tris (pH 8), 15 mM KCl and 0.5 mM Mg^2+^; actual ionic strength also includes ~10 mM NaCl as counterions from the polyions. Turbidity was calculated using UV-Vis absorbance at 500 nm measured by Tecan M1000 Pro microplate reader. Turbidity alone cannot discriminate between aggregates and coacervates solutions.^39^ Therefore, samples were also imaged with a Nikon Eclipse TE200 inverted optical microscope to test the presence of coacervate droplets. Each experiment was repeated at least 3 times. We note that it would very likely be possible to form coacervates from polycation/polyanion pairs that produced aggregates by increasing the solution ionic strength;^27,40,41^ we did not do this here because we wished to hold the buffer conditions constant so as to compare the properties of coacervates formed from different length polyions.

### Salt and pH measurements

pH of coacervate systems (droplet and continuous phase) was estimated using C-SNARF-1 dye emission using 543 nm excitation with confocal microscope. pH of dilute phase was measured by micro pH electrode after centrifugation. Salt concentration of samples were adjusted by concentrated KCl to the desired salt concentrations. Detailed information about experimental setting and calculation is in SI.

### RNA partitioning experiments (fluorescence)

Coacervate samples were prepared according to the coacervate preparation section except the certain volume of water was replaced with volume of labeled RNA(ACCUUGUUCC[Cy3] or GGAACAAGGUAGAGCGAGAU[Cy3]), which was added last, to 0.1 μM final concentration. For dsRNA experiments, equimolar unlabeled complementary RNA sequence (see Supplementary Table 4) was mixed and heated at 95 °C for 2 min. Then, the mixture was left at room temperature for 1 h. Microscope images were taken on the Leica TCS SP5 inverted confocal microscope with exciting wavelength 543 nm. Each experiment was repeated at least three times. From each sample, three images were collected and fluorescence intensity from three droplets per image was measured. Calibration curves were obtained to determine concentrations of labeled RNA in the droplets.

### FRET

We followed the FRET method of Nott et al,^32^ as described in detail in Supplementary Information. RNA 10mer sequences (ACCUUGUUCC[Cy3] and [Cy5]GGAACAAGGU) with and without Cy3 and Cy5 fluorescent labels. RNA were used. Cy3 (donor) was excited at 543 nm and emission collected between 555-625 nm. Cy5 (acceptor) was excited at 633 nm and emission collected between 650-750 nm. Acceptor only, donor only, and FRET samples were used to calculate corrected FRET. Three fluorescence images were obtained for each sample including coacervates and buffer.

### Radioactive tRNA partitioning experiments

Coacervates were prepared by first adding water, then 10mM Tris pH 8.3, 15mM KCl, 0.5mM MgCl_2_, polyanion, then polycation followed by renatured ^32^P labeled tRNA^phe^. This was mixed well and 1μL was added to 10mL of scintillation fluid and counted. To determine quantity of the radiolabeled RNA that remained in the dilute phase, samples were centrifuged for 1 min. at 14k RPM, then 1μL of the resulting dilute phase was pipetted into 10mL of scintillation fluid and counted. Then to calculate % RNA in the coacervate phase, the measured value for the dilute phase was divided by the total. The amount of RNA in the dilute phase was measured in triplicate for each coacervate phase, and in triplicate for the total samples (coacervate + continuous phases). Values in Supplementary Table 7 are averages of three replicates of the dilute phase that were scintillation counted and then averaged.

### In-Line Probing

For the buffer only reactions, 6 kcpm/µL of 5’radiolabled tRNA^phe^ was incubated in 10 mM Tris (pH 8.3) 15 mM KCl, and 0.5 mM MgCl_2_ at 37 °C for up to 48 h. For the polycation-only reactions, tRNA^phe^ was incubated in the above conditions but with 10 mM total (+) charge for each of the polycations. For the polyanion-only reactions, tRNA^phe^ was incubated in the above conditions but with 10 mM total (–) charge for each of the polyanions. For coacervate reactions, tRNA^phe^ was incubated with the above conditions with a 1:1 ratio of + charge to – charge at 10 mM +charge and 10 mM – charge. Reactions were fractionated on 10% denaturing urea polyacrylamide gels at 60 W for 1.5 h before being dried at 70 °C for 1 h. Gels were then exposed on PhosporImager plates overnight before being imaged on a Typhoon scanner.

## Acknowledgements

This work was supported by the NASA Exobiology program grant 80NSSC17K0034. S. C was supported by Future Investigators in NASA Earth and Space Science and Technology (FINESST) under Grant 80NSSC19K1531.

## Author Contributions

C.D.K, P.C.B. and F.P.C designed the systems. F.P.C. performed coacervate formation, partitioning and FRET experiments. S.C. performed salt resistance and pH experiments, and M.M. performed in line probing experiments. C.D.K, P.C.B. and F.P.C wrote the manuscript with contributions from S.C. and M.M.

## Additional Information

Supplementary information is available in the online version of the paper. Reprints and permissions information are available online at www.nature.com/reprints. Correspondence and requests for materials should be addressed to C.D.K and P.C.B.

## Competing interests

The authors declare no competing financial interest.

