## Supplemental Information for "Prebiotically-relevant low polyion multivalency can improve functionality of membraneless compartments"

### Prebiotically-relevant low polyion multivalency provides functional membraneless compartments

**Table of Contents**

Supplementary Materials and Methods…..…………………………………………...... S3-S9

Supplementary In-Line Probing Discussion ……………………………………….... S9-S10

Supplementary Figures……………………………………………………………..... S11-S23

Supplementary Tables ……………………………………………………………… S24-S28

Supplementary References ………………………………………………….…… S29

**Supplementary Materials and Methods**

**Materials**

Poly(L-lysine hydrochloride) (degree of polymerization n = 10, 30 and 100), poly(L-arginine hydrochloride) (n = 10, 30 and 100), poly(L-aspartic acid sodium salt) (n = 10, 30 and 100) and poly(L-glutamic acid sodium salt) (n= 20 and 100) were purchased from Alamanda Polymers and were used without further purification. L-lysine monohydrochloride, L-arginine monohydrochloride, L-aspartic acid sodium salt monohydrate, L-glutamic acid monosodium salt hydrate, adenosine 5'-monophosphate disodium salt (AMP), adenosine 5'-diphosphate disodium salt (ADP) and adenosine 5'-triphosphate disodium salt hydrate (ATP) were purchased from Sigma Aldrich. Poly(L-lysine hydrochloride) (n =5), Poly(L-arginine hydrochloride) (n = 5), Poly(L-aspartic acid hydrochloride) (n = 5), Poly(L-glutamic acid hydrochloride) (n = 5 and 10) were purchased from Genscript. ss RNA 10mer, ss RNA 20 mer were purchased from Sigma Aldrich, details are in SI. SNARF-1, carboxylic acid, acetate, succinimidyl ester was purchased from ThermoFisher Scientific.

**Coacervate preparation**

pH of the stock solutions was fixed to ~8 by adding NaOH or HCl. The order of addition was as follows: water, KCl, $MgCl_2$, Tris, polyanion and polycation. Samples were pipette mixed and transferred to a Corning 96 well special optics plate. Absorbance spectra for each well were recorded from 500 - 600 nm using a Tecan M1000 Pro microplate reader.

**Phase diagram of salt dependent coacervate formation**

Salt dependent coacervate formation is investigated here using turbidity measurement and microscopy confirmation. Same concentration of peptides or ATP, $Mg^{2+}$ and Tris buffers are used. To achieve desired salt concentration, 2 M or 400 mM KCl is used. The pH of all samples is measured by pH electrode and measured as around pH 8.1. Turbidity is calculated as follows:

$$\text{Turbidity (\%)} = 100 - 10\text{\^{}}(2\text{-}Abs_{500nm,\ sample} + Abs_{500nm,\ buffer}) \qquad \text{(Equation 1)}$$

The absorbance at 500 nm of turbid samples was collected using a Tecan M1000 Pro microplate reader. Absorbance values at 500 nm were pathlength corrected according to the application notes by Thermo Fisher Scientific Inc.[1] Turbidity curves as a function of KCl concentration were fit to a the Hill equation (equation 2), using weighted fitting by IgorPro Version 6.37 software. The relative error was defined as the square root of the sum of the square of 10% of experimental value (to account for systematic errors) and the square of the standard deviation for multiple measurements. The fitting parameter $K_{1/2}$ is the concentration of $K^+$ at the one-half y-maximum, which indicates the transition salt concentration of coacervate formation in Fig. 3 (Supplementary Table 1). The fitting parameter $T_{max}$ corresponds to the maximum turbidity at 0 mM KCl, which was determined from the fit. The fitted value of *n* ranged from 4.7 ~ 17.6 according to the steepness of the curve, but since it lacked physical meaning it was not reported. In deriving Equation 2, the minimum value of the turbidity was set to zero, which is consistent with all of the data.

$$T = T_{max}\left[\frac{1}{([\mathrm{K}^+]/K_{1/2})^n + 1}\right] \qquad \text{(Equation 2)}$$

**Measurement of local pH in coacervate droplets**

To determine pH, we used 5-(and-6)-carboxy SNARF®-1 (C-SNARF-1), which is a widely used ratiometric pH indicators to measure local pH in cells[2] or polymer microspheres,[3] on charged surface nanoparticles[4]. C-SNARF-1 is especially appealing as pH indicator to measure local pH of coacervate droplets, since the ratiometric properties of C-SNARF-1 are not significantly dependent on its concentration or on the ionic strength of the surrounding aqueous media.[2] The ratio of fluorescence intensity of dual emission peaks was shown to be invariant upon photobleaching.[2] One common approach to determining local pH of biological samples has been using equation (3), assuming the pKa of SNARF is a fixed value between pH 7.1 ~ 7.5[5].

$$pH = pK_A - \log\left(\frac{R-R_B}{R_A-R} \times \frac{F_{B(\lambda 2)}}{F_{A(\lambda 2)}}\right) \quad \text{(Equation 3)}$$

This is based on the binding formula using the ratio of the emission (R) at 580 - 590 nm ($\lambda 1$) and 625 - 635 nm ($\lambda 2$). The subscript A and B are the acidic end point and basic endpoints, respectively. The fluorescence emission (F) values at $\lambda 2$ for the basic and the acidic end point are required for this formula. Supplementary Equation (3) can be modified into linear relationship between pH and $\boldsymbol{LogE}$ as in Supplementary equation (4). This allow us to avoid using predetermined fixed pKa values of C-SNARF-1, and consequently, predict the apparent pH of dilute phase well. This method has been used in analytical assessment of this pH probe and measuring pH gradients in microfluidic devices.[6,7]

$$pH = pKa - \log\left(\frac{F_{B(\lambda 1)}}{F_{A(\lambda 2)}}\right) - \log\left(\frac{R-R_B}{R_A-R}\right) = \alpha - \beta \log\left(\frac{R-R_B}{R_A-R}\right) = \alpha - \beta\ \boldsymbol{LogE} \quad \text{(Equation 4)}$$

α and β are experimentally determined values by linear fitting using Supplementary equation (4) to the curve of pH versus $\boldsymbol{LogE}$ as in Supplementary Fig. 4 (A). The detailed fitting method and fitting parameters are described in Supplementary Fig. 4.

C-SNARF-1 was purchased from Thermo Fisher Scientific (CAS Number 126208-12-6). C-SNARF-1 stock solution was made as 1 mM in DMSO and mixed with coacervate solutions to a final concentration of 25 µM. For the calibration curves in Supplementary Fig. 3, emission of SNARF from 560 nm to 700 nm was measured by a Jobin Yvon Horiba FL3-21 fluorimeter with 5 nm slit size, 5 average scan and 543 nm excitation, and the pH of solution is adjusted by addition of NaOH or HCl solutions and measured by Mettler Toledo Ultra Micro ISM electrode. The calibration curve of SNARF emission ratio is achieved in phosphate buffer. Next, C-SNARF-1 emission from coacervate droplets in/ out is collected by lambda scan using Olympus Fluoview 1000 Confocal Microscope simultaneously. We used 543 nm laser with varied laser intensity and gain to prevent the saturation of fluorescence emission; 35- 50% laser intensity and 500 – 650 V gain. RP 20/80 filter was utilized, and lambda scan was conducted with 5 nm step size and 10 nm bandwidth. The lambda scan of coacervate samples without C-SNARF-1 is performed for the baseline correction with the same setting of confocal microscope. Three or more sets of samples were prepared. 5 ROIs per image were utilized to calculate local pH of coacervate droplets, and three images were taken from each sample. Additionally, pH of supernatant phase is measured by micro pH probe after 2 hours of equilibration and 15 mins of centrifugation. We chose to use the calibration curve of SNARF in phosphate buffer since the apparent pH of dilute phase were estimated very close to the pH of dilute phase measured by pH electrode. The variation of apparent pH of dilute phases by fitting curves is plotted as compared to the dilute phase pH values measured by pH electrode in Supplementary Fig. 4.

**RNA Partitioning experiments (continuous phase)**

Bulk fluorescence measurements were made using Fluorolog 3-21 fluorimeter with FluorEssence software and a Wavelength Electronics temperature controller. Coacervate and continuous phases separated by centrifugation and fluorescence in continuous phase was measured using fluorimeter. Calibration curves of known concentration of labelled RNA were prepared by using fluorimeter and used to determine concentration in continuous phase. Partitioning coefficient was calculated by dividing concentration in droplet phase by continuous phase.

**FRET**

We used same method applied in a previous study.[8] We used Cy3 and Cy5 as fluorophores a FRET pair. Cy3 (donor) is excited at 543 and emission is collected between 555-625 nm. The Cy5 (acceptor) is excited at 633 nm and emission is collected between 650nm-750 nm. Three fluorescence images were obtained for each sample including coacervates and buffer.

Those fluorescence channels correspond to following:

$DD_{obs}$= observed donor emission after donor excitation

$DA_{obs}$= observed acceptor emission after donor excitation (FRET)

$AA_{obs}$= observed acceptor emission after acceptor excitation.

To correct for overlap of emission and absorbance of acceptor and donor dyes, we used samples containing only donor or acceptor fluorophores with same parameters. Again, three fluorescence images were recorded including DDdonor /DAdonor or DAaccept/AAaccept. Observed emission is either for donor only sample or acceptor only sample, and abbreviation properly is replaced with donor or accept instead of obs accordingly. Image processing and analysis were accomplished by using Fiji. The corrected FRET, $E_{CT,}$ is calculated as:

$$E_{CT} = \frac{DA}{DA + DD} = \frac{DA_{obs} - \alpha \cdot DD_{obs} - \beta \cdot AA_{obs}}{DD_{obs} + (DA_{obs} - \alpha \cdot DD_{obs} - \beta \cdot AA_{obs})}$$

Correction terms are calculated as follows:

$$\alpha = \frac{DA_{donor}}{DD_{donor}} \qquad and \qquad \beta = \frac{DA_{accept}}{AA_{accept}}$$

For each system, five droplets or region of interest (ROI) per image were chosen from three images and each measurement was repeated with three different samples. Mean intensity was calculated with Fiji-ImageJ. Mean intensities were used to calculate average correction factors ($\alpha$ $and$ $\beta$) and used to calculate corrected FRET.

**In vitro transcription**

The tRNA$^{phe}$ was *in vitro* transcribed by T7 RNA polymerase with the following conditions: 40 mM Tris (pH 7.5), 3mM each NTP, 25 mM $MgCl_2$, 2 mM DTT, 0.14 µM template DNA and 0.14 µM T7 promoter DNA, and 6% by volume T7 Polymerase. Template DNA and T7 promoter DNA were mixed with 10 mM NaCl and 1X TBE before being renatured at 90 $^oC$ for 1.5 min then cooled to room temperature for 5 min. This mixture was added to the rest of the mixture and finally 6% by volume T7 polymerase was added before being incubated at 37 $^oC$ for 4 h. The transcription reaction was quenched by addition of 2X formamide loading dye containing 10 mM EDTA, 90% formamide 0.025% Bromophenol Blue. This was then loaded onto a 10% denaturing urea polyacrylamide gel and fractionated at 25 W for ~1.5 h before the RNA was visualized by UV shadowing and cut out. The gel slice was then crushed and soaked overnight in a buffer that contained 250 mM Tris (pH 8.0), 250 mM ethylenediaminetetraacetic acid (EDTA), and 250 mM NaCl (TEN 250) before being ethanol precipitated.

**SAP/Kinase**

Before incubation with Shrimp Alkaline Phosphatase (SAP) to remove the 5'-triphosphate of the RNA, the RNA was renatured at 95 $^oC$ for 1 min, then allowed to return to room temperature. Purified in vitro transcribed tRNA$^{phe}$ was incubated with 3 U SAP in 1X Cutsmart buffer at 37

°C for 1 h. Next, SAP was heat inactivated at 65 °C for 5 minutes. Kinase reaction conditions: 4.9 μM $tRNA^{phe}$, 10 U of T4 Polynucleotide Kinase, 1X PNK buffer, 10% DMSO, 2.5 μM $\gamma$-$^{32}$P ATP. Kinase reaction was incubated at 37 °C for 1.5 h before being quenched with 2X formamide loading dye and loaded onto a 10% denaturing urea polyacrylamide gel fractionated at 20 W for 1 h. The gel was imaged using X-ray film and subsequently the RNA band was excised from the gel placed in TEN 250 buffer and crush and soaked overnight at 4 °C. The next day it was ethanol precipitated and scintillation counted.

**In-Line Probing (ILP) control discussion**

To assess its effect on the folding state of the tRNA, ILP was performed with each length of Lys homopolymer. All three Lys homopolymers (n=10, 30, 100) led to partial unfolding of the tRNA (Supplementary Fig. 11). The acceptor stem is particularly reactive, especially when incubated with the $(Lys)_{10}$ homopolymer. Reactivity at the 5'-end of this stem is unlikely to be due to the reaction going beyond a single reaction because of the low $Mg^{2+}$ ion concentration of 0.5mM and because generally after 24 h, the RNA is in the single-hit regime as judged by the extensive starting material remaining. $(Lys)_{10}$ also leads to unfolding of the acceptor and D-stems, with general unfolding elsewhere. In contrast, although $(Lys)_{30}$ and $(Lys)_{100}$ also led to the acceptor stem being unfolded, there is not as intense reactivity in the D-stem, AC-stem and TC nucleotides in these reactions.

Additionally, ILP was performed with all of the anions individually (Supplementary Fig. 12). With short aspartate polymers n=5, 10, 30, the tRNA folds natively, as judged by comparison to the buffer only in-line probing. However, overall ILP reactivity decreases with increased polymer length. Loss of ILP reactivity is presumably due to the increased multivalency of the polymer increasing its ability to chelate $Mg^{2+}$ ions, which are required for the ILP reaction. With $(Asp)_{100}$ alone, however, there is unfolding of the acceptor stem and gain of reactivity in the variable loop. This may be due to a specific two hydrogen-bond interaction of Asp's carboxylates with the Watson-Crick faces of guanine residues that is weaker at shorter polymer lengths. When incubated with ATP-only, there is essentially no reactivity, due to the high affinity of ATP for $Mg^{2+}$ ions leading to strong chelation preventing the ILP reaction from proceeding.[9] Consistent with this notion, if the ATP is first mixed in a 1:1 molar ratio with $Mg^{2+}$ ions before using the standard ILP conditions (10mM Tris, pH 8.3, 15mM KCl, 0.5mM $MgCl_2$) to saturate the phosphates, ILP reactivity is regained and the native secondary structure pattern is seen.

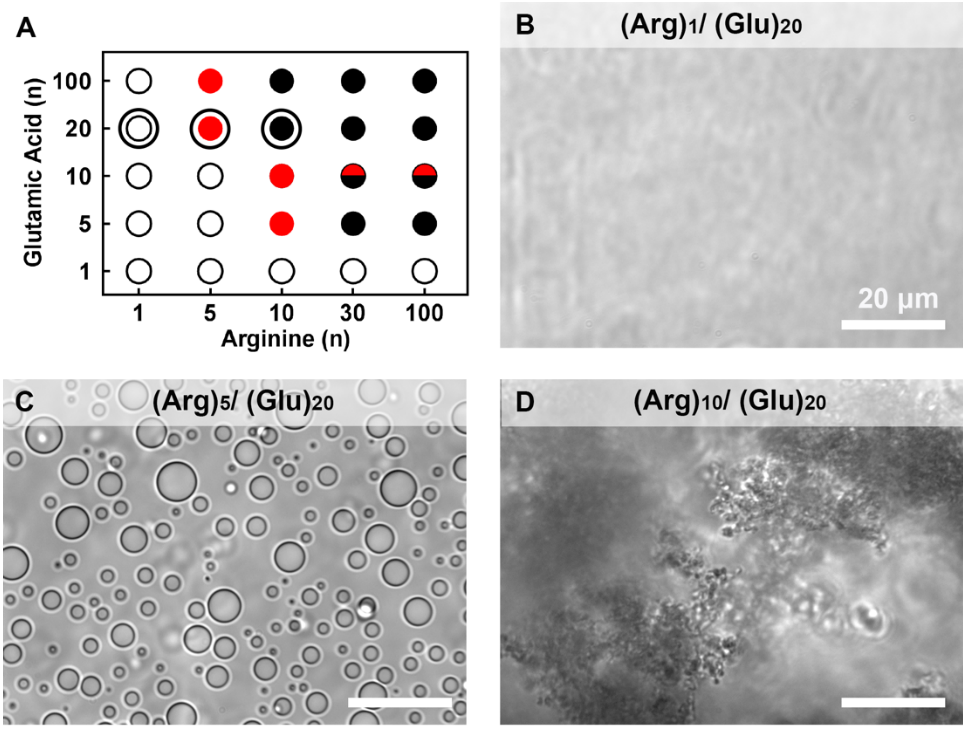


**Supplementary Figure 1.** Interaction of glutamic acid with arginine. (**A**) Summary of conditions tested. (**B**)- (**D**)The subsequent panels highlight interaction of $(Glu)_{20}$ with different length of $(Arg)_n$ (n = 1, 5 and 10, respectively) leading to **(B)** uniform solution, **(C)** coacervate and **(D)** aggregates. Shown are microscope images of uniform solution ( ○ ), coacervates ( ● ) and aggregates ( ● ), selected and highlighted in graph **(A)**.

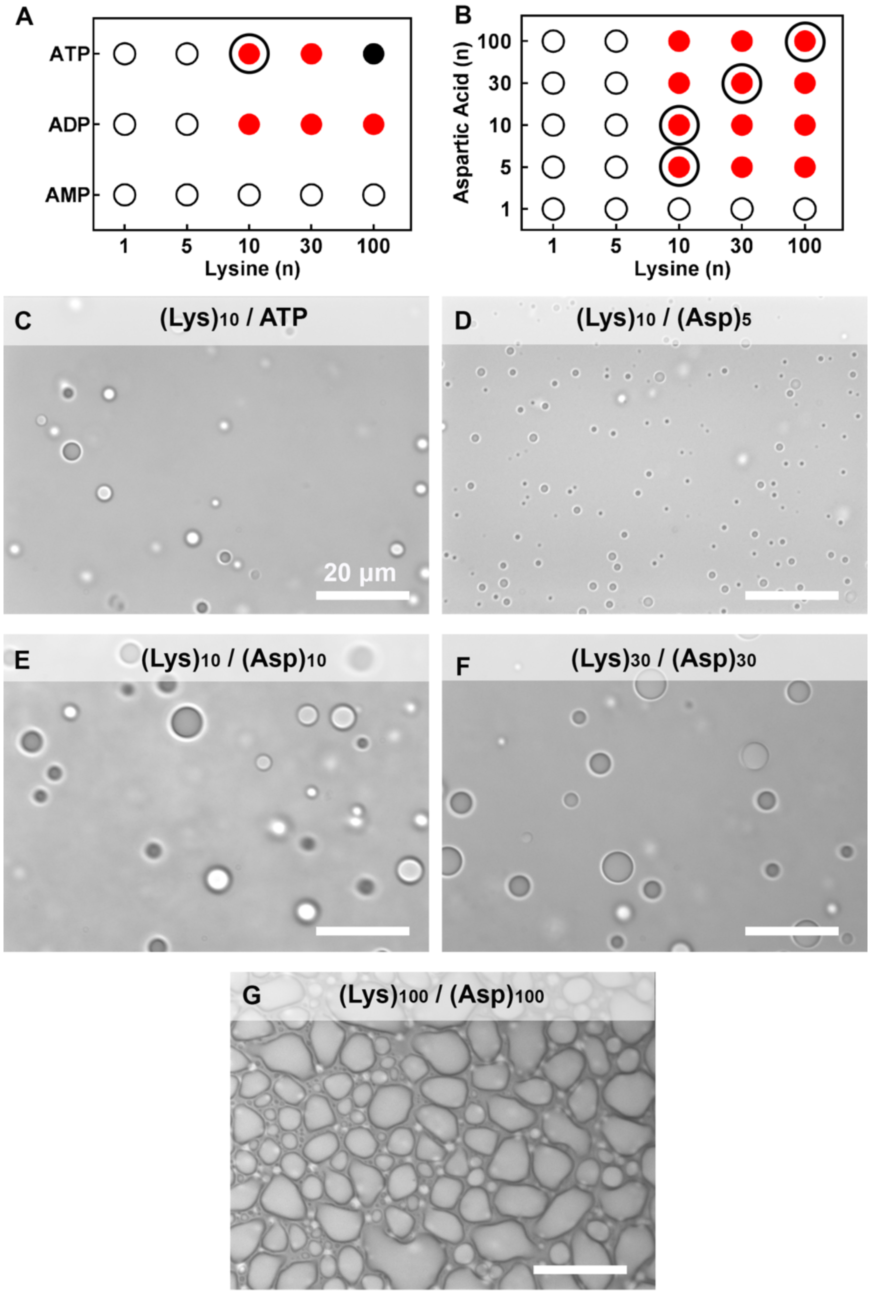


**Supplementary Figure 2.** Coacervate systems chosen for further experiments. Interactions of $(Lys)_n$ with **(A)** nucleotides (AMP, ADP and ATP) and **(B)** $(Asp)_n$ (n= 1, 5, 10, 30 and 100). Specific polyion pairs used to compare across a range of multivalency are circled in panels A and B. Symbols indicate uniform solution ( ○ ), coacervates ( ●) and aggregates ( ● ). Coacervate images are shown for **(C)** $(Lys)_{10}$/ATP, **(D)** $(Lys)_{10}/(Asp)_5$, **(E)** $(Lys)_{10}/(Asp)_{10}$, **(F)** $(Lys)_{30}/(Asp)_{30}$ and **(G)** $(Lys)_{100}/(Asp)_{100}$ pairs.

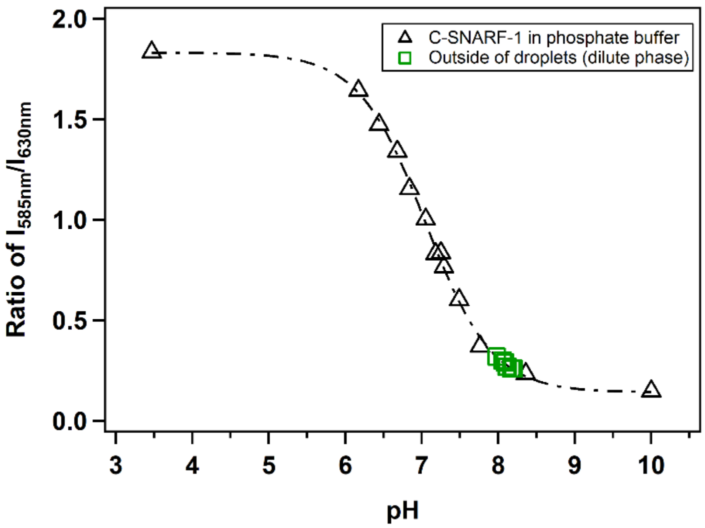


**Supplementary Figure 3.** Calibration curves of SNARF-1 with emission ratio of SNARF-1 in coacervate droplets. Ratio of intensity at 585 nm and 630 nm at various pHs. Black open triangle **(**Δ**)** is ratio of C-SNARF-1 (25 μM) in phosphate buffer (10 mM). Calibration is fitted with the sigmoid curve. Open green square (□): R values collected from coacervate droplets outside (dilute phase) by confocal microscope as a function of pH measured from pH electrode. These data points of pH of dilute phase measured by pH versus ratio (□) are well overlapped with phosphate buffer curve **(**Δ**)**.

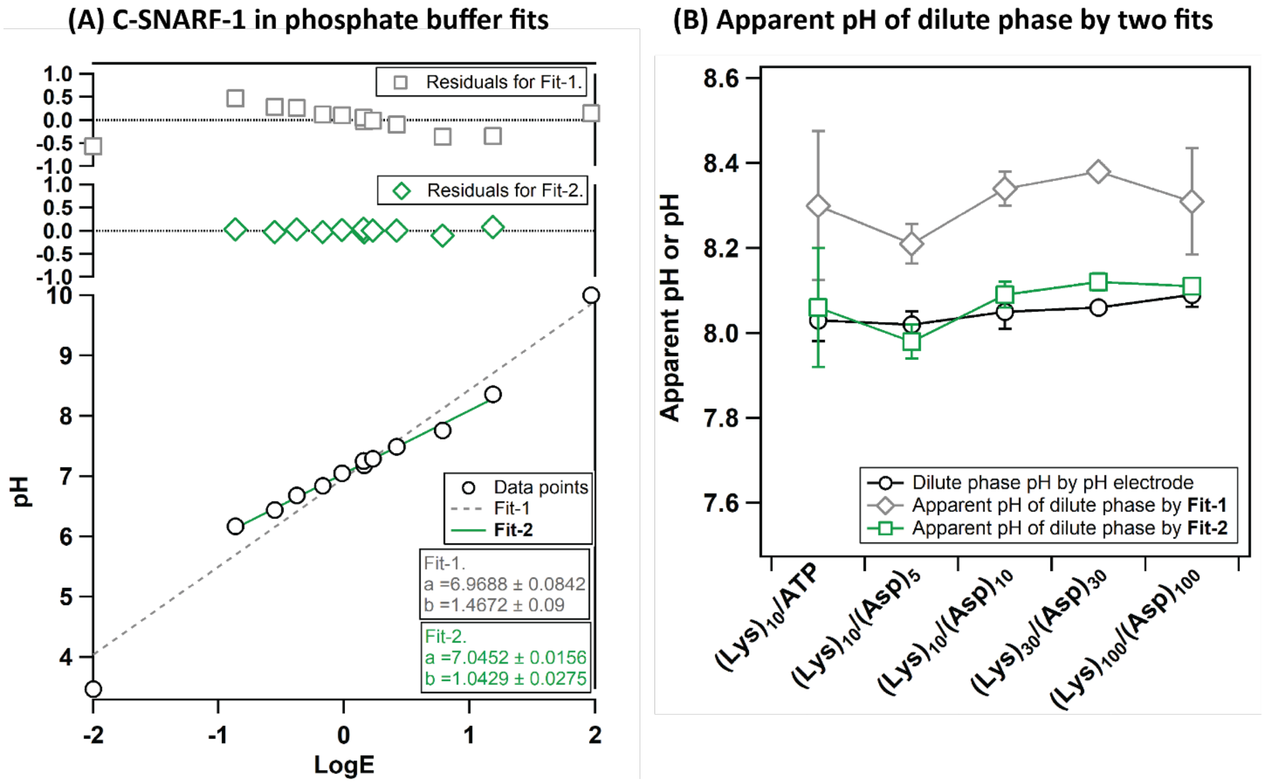


**Supplementary Figure 4**. (A) Calibration of C-SNARF-1 in phosphate buffer. Linear calibration curves and residuals are based on Supplementary equation (4). Linear fitting is performed using IgorPro Version 6.37 software and its fitting equation is y = a + b x (a and b are fitting parameters), whereas a = - α and b = β, y = pH by pH electrode and x = **Log E** from Supplementary equation (4). Fit-1 is the linear fit including all data points and Fit-2 is the linear fit excluding two end points for the better fit. These two data points are excluded since they are not in the transition range from Supplementary Fig. 3. (B). The comparison of apparent pH values of dilute phase to the pH measured by pH electrode. Apparent pH values of dilute phase are estimated using two different fitting equations from (A). We confirmed the Fit-2 from (A) can estimate the apparent pH of dilute phase closer to the pH values measured by pH electrode (□) than Fit-1 (◊). Therefore, we use this linear equation to estimate the apparent pH of coacervate droplets in Fig. 3 (E).

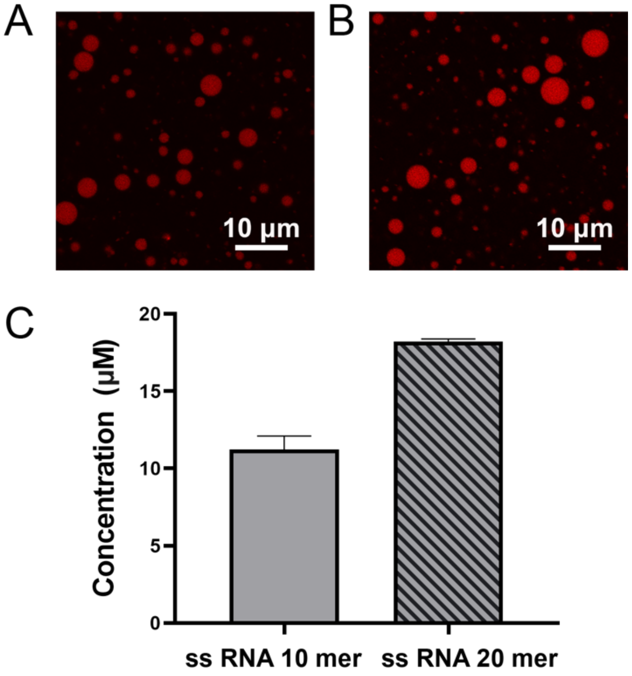


**Supplementary Figure 5.** Partitioning of fluorescently labelled ssRNA 10 and 20mers in of $(Lys)_{100}/(Asp)_{100}$ coacervate pair at 1.2:1 +:– charge ratio. **(A, B)** Fluorescence image of Cy3 labelled ssRNA **(A)** 10mer and **(B)** ssRNA 20mer partitioned in coacervate droplets of $(Lys)_{100}/(Asp)_{100}$. **(C)** Calculated concentration of ssRNA 10 and 20mer in the droplets. Labelled ssRNA was added to a final concentration of 0.1 μM.

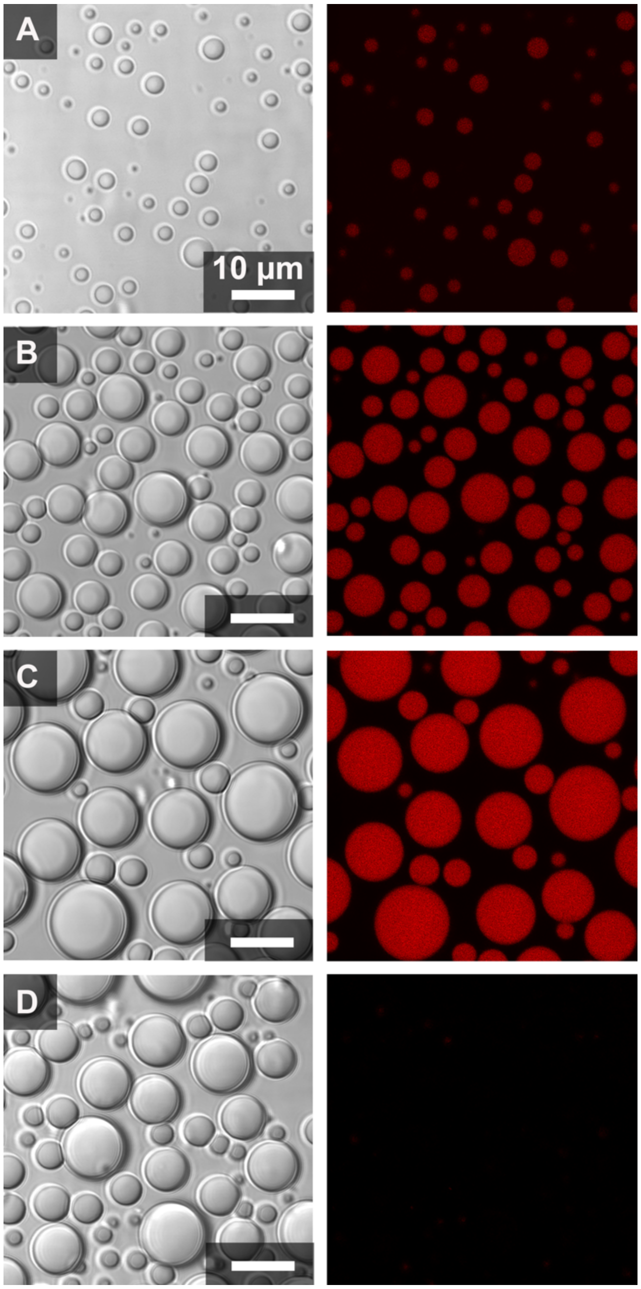


**Supplementary Figure 6** Optical microscope images showing transmitted light (left) and fluorescent channel (right) of coacervate systems. Fluorescently labelled ssRNA 10mer partitioning in **(A)** $(Lys)_{10}$ /$(Asp)_{5}$, **(B)** $(Lys)_{10}$ /$(Asp)_{10}$, **(C)** $(Lys)_{30}$ /$(Asp)_{30,}$ and **(D)** $(Lys)_{100}$ /$(Asp)_{100}$ coacervate pairs. Note that laser intensity was optimized separately for each sample; quantification based on calibration curves is shown in Figure 4C.

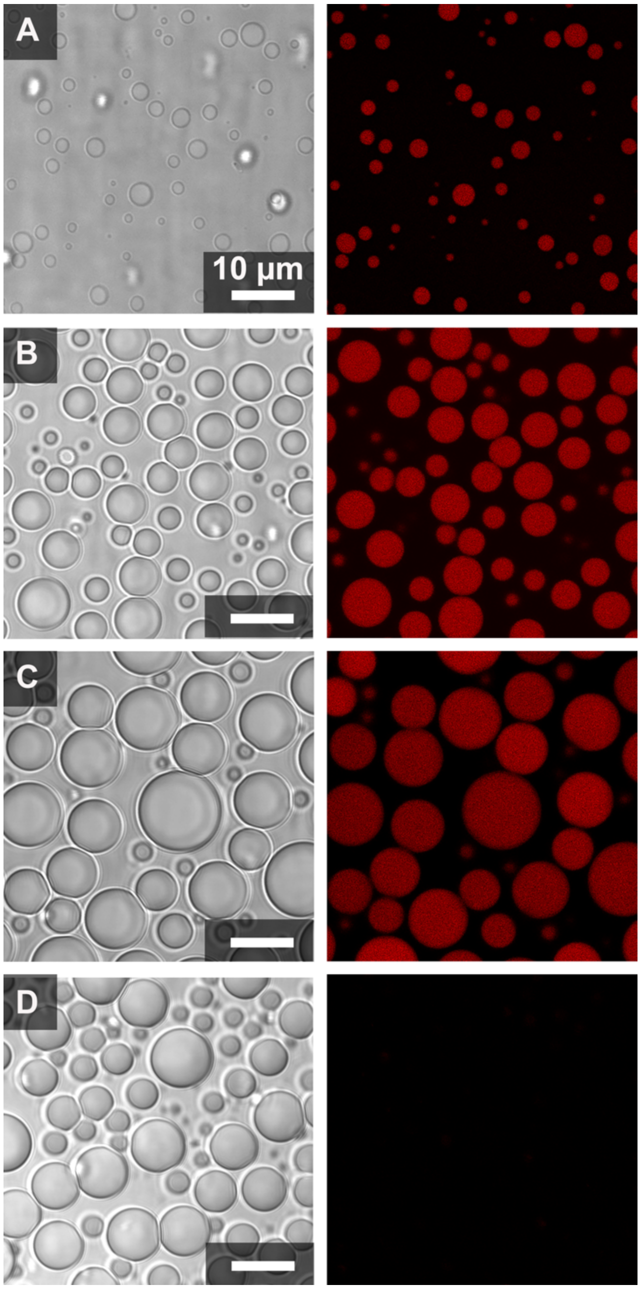


**Supplementary Figure 7** Optical microscope images showing transmitted light (left) and fluorescent channel (right) of coacervate systems. Fluorescently labelled ssRNA 20mer partitioning in **(A)** $(Lys)_{10}$ /$(Asp)_{5}$, **(B)** $(Lys)_{10}$ /$(Asp)_{10}$, **(C)** $(Lys)_{30}$ /$(Asp)_{30}$, and **(D)** $(Lys)_{100}$ /$(Asp)_{100}$ coacervate pairs. Note that laser intensity was optimized separately for each sample; quantification based on calibration curves is shown in Figure 4C.

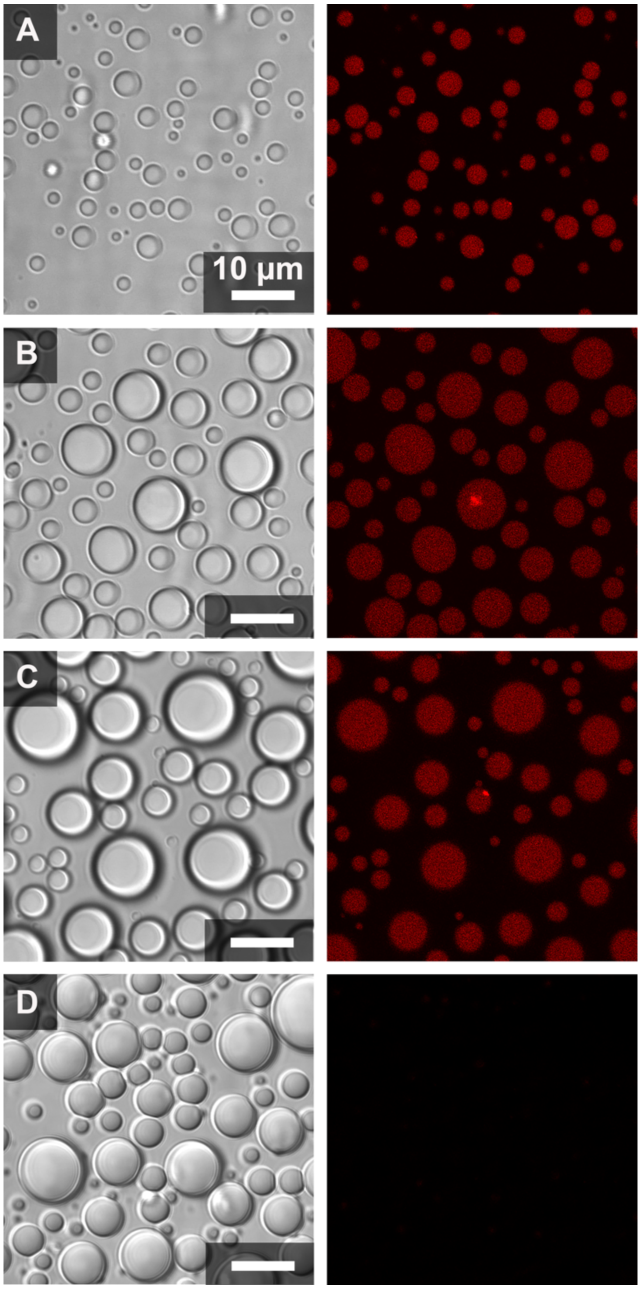


**Supplementary Figure 8.** Optical microscope images showing transmitted light (left) and fluorescent channel (right) of coacervate systems. Fluorescently labelled dsRNA 10mer partitioning in **(A)** $(Lys)_{10}/(Asp)_5$, **(B)** $(Lys)_{10}/(Asp)_{10}$, **(C)** $(Lys)_{30}/(Asp)_{30}$, and **(D)** $(Lys)_{100}/(Asp)_{100}$ coacervate pairs. Note that laser intensity was optimized separately for each sample; quantification based on calibration curves is shown in Figure 4C.

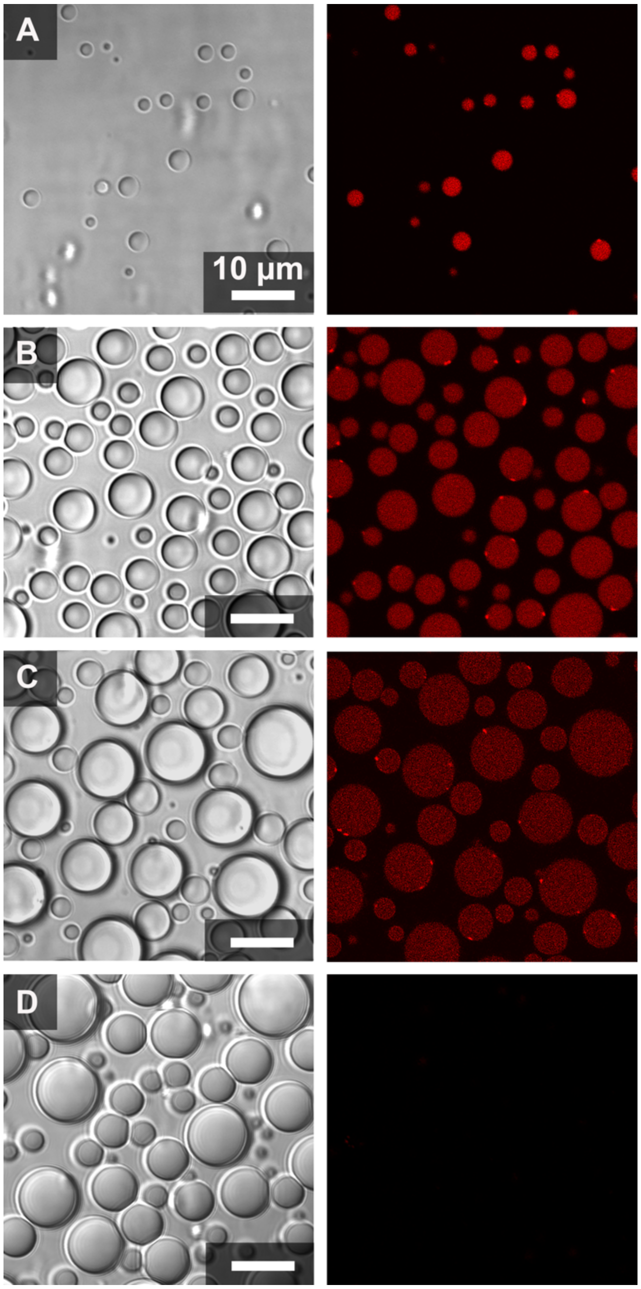


**Supplementary Figure 9.** Optical microscope images showing transmitted light (left) and fluorescent channel (right) of coacervate systems. Fluorescently labelled dsRNA 20mer partitioning in **(A)** $(Lys)_{10}$ /$(Asp)_5$, **(B)** $(Lys)_{10}$ /$(Asp)_{10}$, **(C)** $(Lys)_{30}$ /$(Asp)_{30}$, and **(D)** $(Lys)_{100}$ /$(Asp)_{100}$ coacervate pairs. Note that laser intensity was optimized separately for each sample; quantification based on calibration curves is shown in Figure 4C.

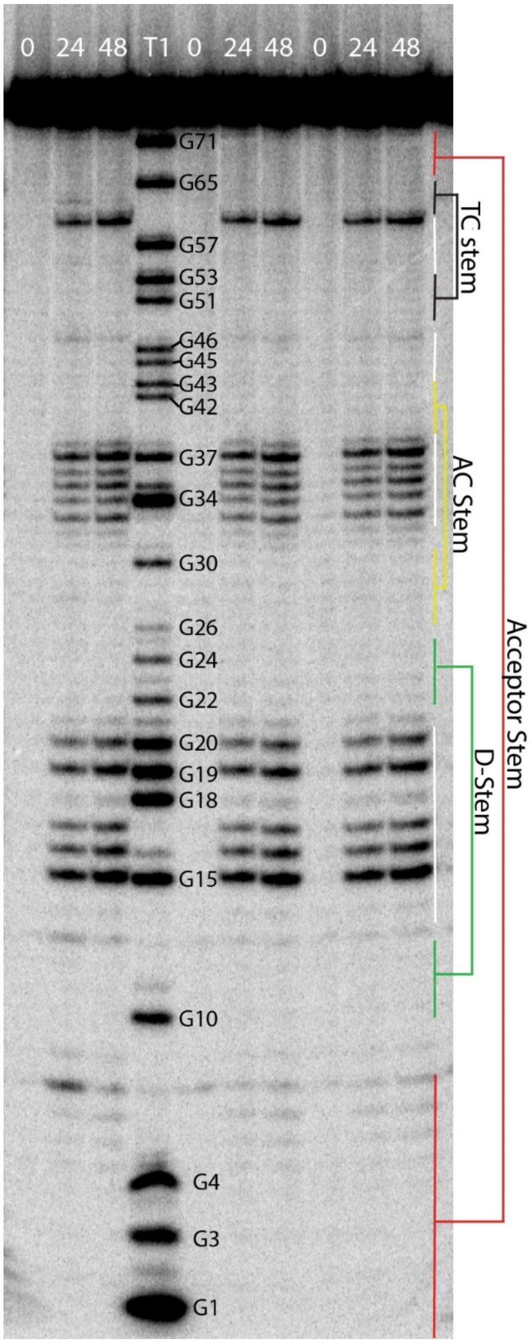


**Supplementary Figure 10.** Gel depicting ILP of tRNA$^{phe}$ in buffer. ILP was performed in triplicate with time-points taken at 0, 24 and 48 h. Buffer conditions were 15mM KCl, 0.5mM $MgCl_2$, 10 mM Tris (pH 8.3).

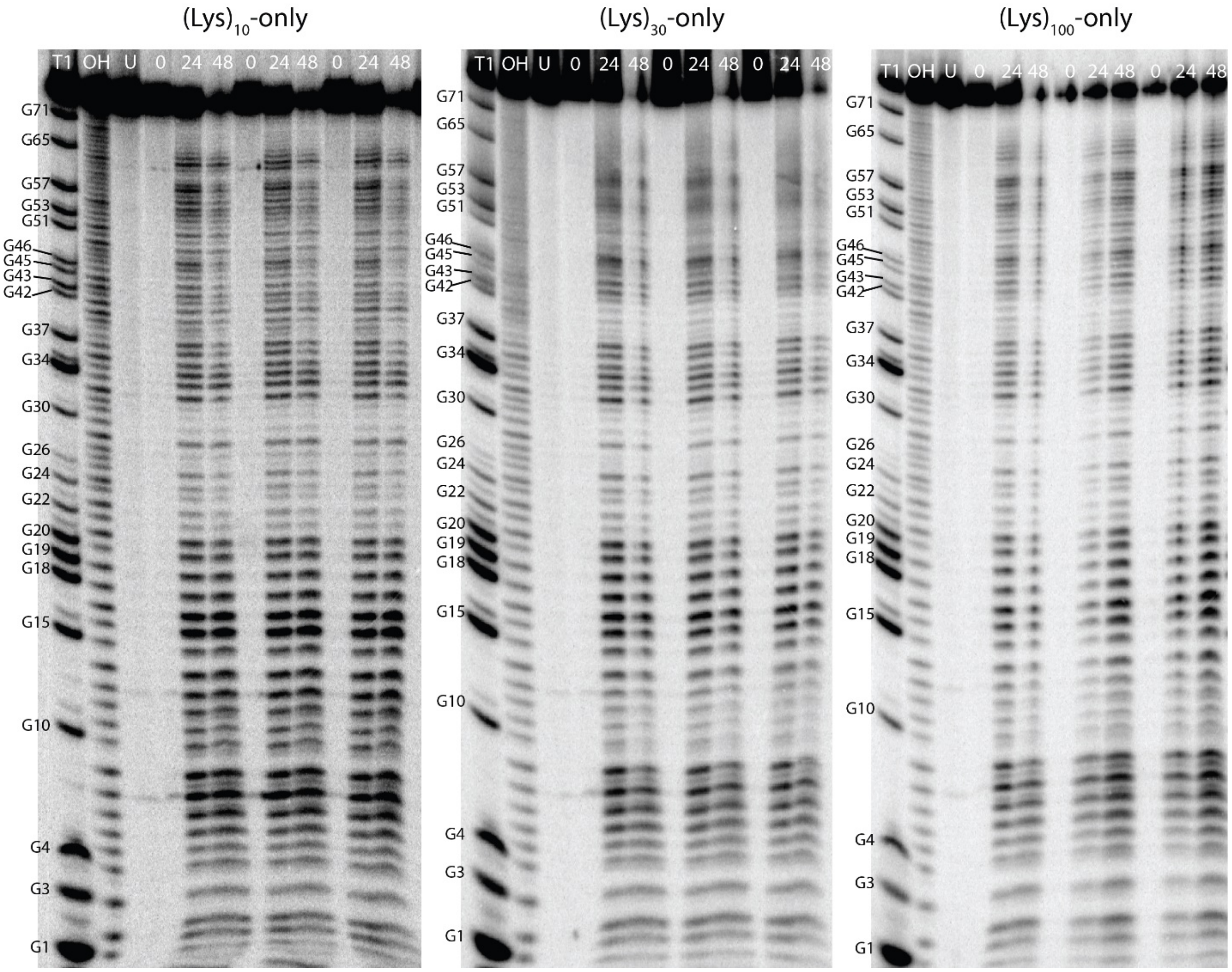


**Supplementary Figure 11.** Gels depicting ILP of tRNA$^{phe}$ incubated with polycations. ILP was performed in triplicate in the presence of the polycations tested (i.e. no coacervates). Under all of these conditions, the tRNA is partially unfolded.

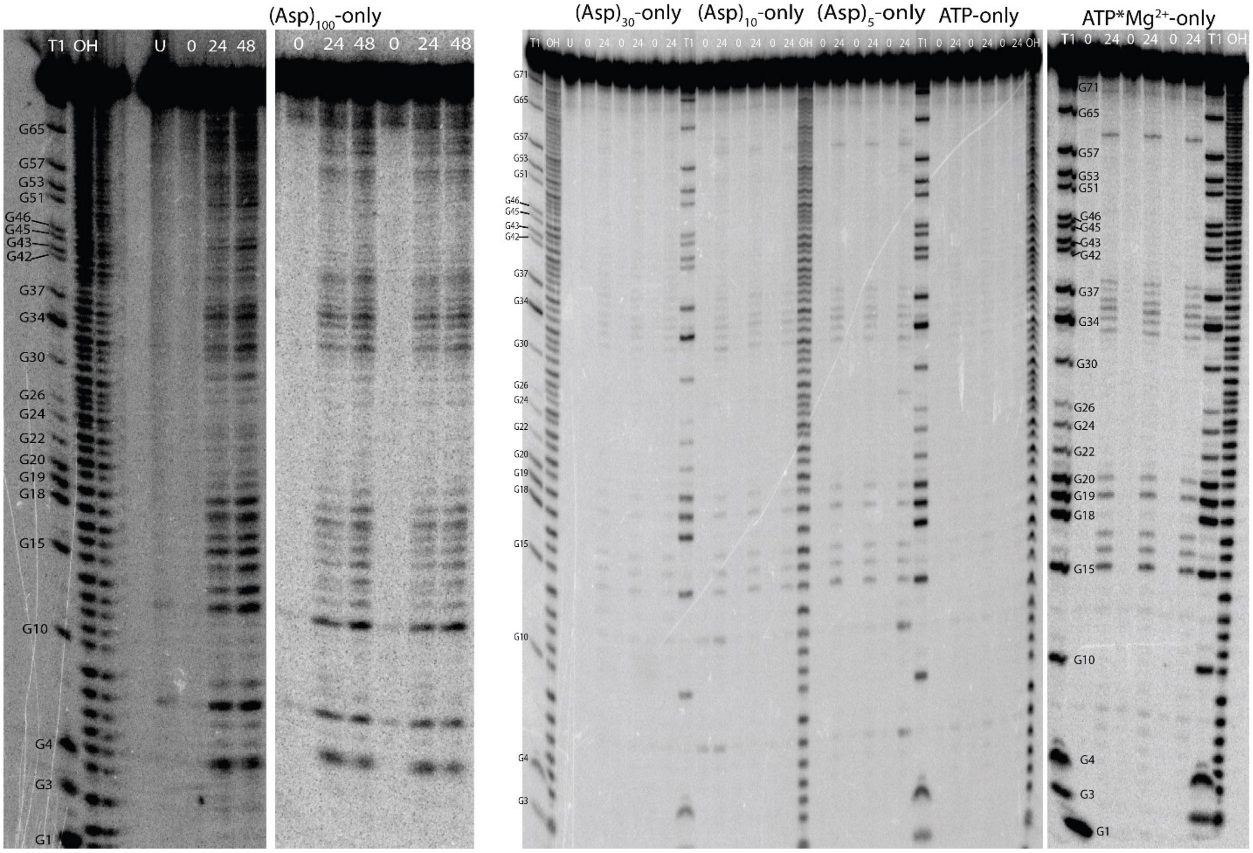


**Supplementary Figure 12.** Gels depicting ILP of tRNA$^{phe}$ incubated with polyanions. ILP was performed in triplicate in the presence of the individual polyanions tested (i.e. no coacervates). For the shorter polyanions, the tRNA remains natively folded and has relatively low ILP reactivity, likely due to partial sequestration of $Mg^{2+}$ ions by the polyanions. However, the $(Asp)_{100}$-only condition shows unfolding comparable to that in coacervates.

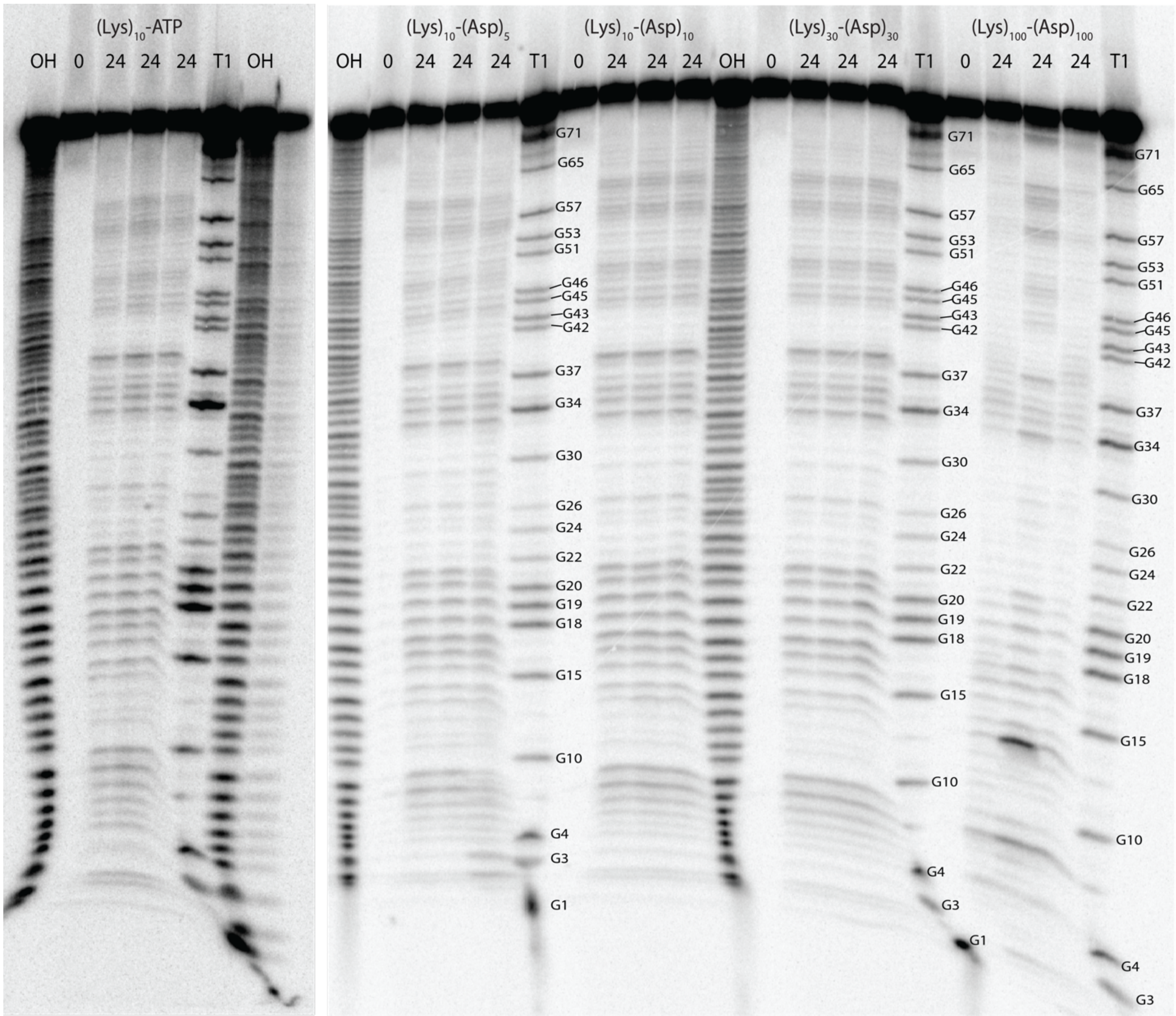


**Supplementary Figure 13.** Gel depicting ILP traces in each of the coacervate conditions tested. ILP for each condition was performed in triplicate. The ILP reactivities were similar between all of the coacervate conditions, revealing unfolding of the acceptor stem, suggesting a common mechanism of unfolding.

**Supplementary Table 1. Transition salt concentration of coacervates (K½) from the fitting in Fig. 3 based on equation (2).**

| Cation | Anion | $K_{½}$, mM | Cation | Anion | $K_{½}$, mM |
|---|---|---|---|---|---|
| $(Lys)_{10}$ | $(Asp)_{5}$ | 20 ± 10 | $(Arg)_{5}$ | $(Asp)_{10}$ | 50 ± 5 |
| $(Lys)_{10}$ | $(Asp)_{10}$ | 230 ± 10 | $(Arg)_{10}$ | $(Asp)_{5}$ | 250 ± 20 |
| $(Lys)_{30}$ | $(Asp)_{30}$ | 440 ± 10 | $(Arg)_{10}$ | $(Asp)_{10}$ | 1320 ± 90 |
| $(Lys)_{100}$ | $(Asp)_{100}$ | 1110 ± 40 | $(Arg)_{10}$ | $(Glu)_{5}$ | 98 ± 3 |
| $(Lys)_{10}$ | $(Glu)_{10}$ | 100 ± 8 | $(Arg)_{10}$ | $(Glu)_{10}$ | 572 ± 8 |
| $(Lys)_{10}$ | ADP | 42 ± 2 | $(Arg)_{10}$ | AMP | 64 ± 8 |
| $(Lys)_{10}$ | ATP | 110 ± 10 | $(Arg)_{10}$ | ADP | 460 ± 20 |
| | | | $(Arg)_{10}$ | ATP | 700 ± 40 |

**Supplementary Table 2. Partitioning of ssRNA 10mer and ssRNA 20mer.**

| Pair of coacervates | RNA 10mer | | | RNA 20mer | | |
|---|---|---|---|---|---|---|
| | Concentration in droplets (μM) | Concentration in continuous phase (μM) | Partitioning coefficient, K | Concentration in droplets (μM) | Concentration in continuous phase (μM) | Partitioning coefficient, K |
| $(Lys)_{10}$/ATP | 42.7 ± 7.1 | 0.0080 ±0.0025 | 5300 ± 1900 | 14.5 ± 2.4 [a] | 0.0038 ±0.00053 | 3600 ± 830 |
| $(Lys)_{10}/(Asp)_5$ | 11.1 ± 0.35 | 0.058 ±0.0095 | 190 ± 32 | 50.7 ± 5.8 | 0.019 ±0.0066 | 2600 ± 980 |
| $(Lys)_{10}/(Asp)_{10}$ | 10.8 ± 2.1 | 0.066 ±0.0069 | 160 ± 36 | 29.4 ± 2.4 | 0.011 ±0.0029 | 2700 ± 740 |
| $(Lys)_{30}/(Asp)_{30}$ | 11.4 ±4.4 | 0.028 ±0.0047 | 400 ± 170 | 11.9 ± 0.37 | 0.028 ±0.013 | 430 ± 24 |
| $(Lys)_{100}/(Asp)_{100}$ | NA [b] | 0.068 ±0.016 | NA [b] | NA [b] | 0.069 ± 0.078 | NA [b] |

[a] This number reflects the concentration of dye in the droplets without aggregation. Final concentration of added RNA is 0.1 μM.
[b] Concentration in the coacervate phase was too low to determine for these samples and consequently a partitioning value could not be calculated.

**Supplementary Table 3. Partitioning of dsRNA 10mer and dsRNA 20mer.**

| Pair of coacervates | RNA 10mer | | | RNA 20mer | | |
|---|---|---|---|---|---|---|
| | Concentration in droplets (μM) | Concentration in continuous phase (μM) | Partitioning coefficient, K | Concentration in droplets (μM) | Concentration in continuous phase (μM) | Partitioning coefficient, K |
| $(Lys)_{10}$/ATP | 13.0 ± 0.71 [a] | 0.090 ±0.011 | 140 ± 19 | 10.5 ± 0.63 [a] | 0.077 ±0.016 | 140 ± 30 |
| $(Lys)_{10}/(Asp)_5$ | 39.9 ± 2.6 | 0.011 ±0.0057 | 3600 ± 1900 | 44.2 ± 0.31 | 0.019 ±0.0059 | 2300 ± 720 |
| $(Lys)_{10}/(Asp)_{10}$ | 18.3 ± 0.84 | 0.067 ±0.0087 | 270 ± 38 | 7.8 ± 1.7 | 0.093 ±0.010 | 84 ± 20 |
| $(Lys)_{30}/(Asp)_{30}$ | 6.3 ± 0.26 | 0.0094 ±0.0048 | 700 ± 340 | 1.4 ± 0.082 | 0.060 ±0.0077 | 23 ± 3 |
| $(Lys)_{100}/(Asp)_{100}$ | NA [b] | 0.097 ±0.012 | NA [b] | NA [b] | 0.091 ± 0.013 | NA [b] |

[a] This number reflects the concentration of dye in the droplets without aggregation. Final concentration of added RNA is 0.1 μM.
[b] Concentration in the coacervate phase was too low to determine for these samples and consequently a partitioning value could not be calculated.

**Supplementary Table 4. RNA sequences used for experiments and corresponding names.**

| Oligo name | Length (nt) | Sequences (5' to 3') |
|---|---|---|
| ssRNA 10mer – Cy3 | 10 | ACCUUGUUCC[Cy3] |
| ssRNA 10mer – Cy5 | 10 | [Cy5]GGAACAAGGU |
| ssRNA 10mer | 10 | ACCUUGUUCC |
| ssRNA 10mer | 10 | GGAACAAGGU |
| ssRNA 20mer– Cy3 | 20 | AUCUCGCUCUACCUUGUUCC [Cy3] |
| ssRNA 20mer– Cy5 | 20 | [Cy5]GGAACAAGGUAGAGCGAGAU |
| ssRNA 20mer | 20 | AUCUCGCUCUACCUUGUUCC |
| ssRNA 20mer | 20 | GGAACAAGGUAGAGCGAGAU |

**Supplementary Table 5**. **RNA sequences including sense and antisense used for FRET experiments.**

| Oligo name | Length (bp) | Sequence (sense) (5' to 3') | Sequence (antisense) (5' to 3') |
|---|---|---|---|
| dsRNA 10mer - Donor | 10 | ACCUUGUUCC[Cy3] | GGAACAAGGU |
| dsRNA 10mer - Acceptor | 10 | [Cy5]GGAACAAGGU | ACCUUGUUCC |
| dsRNA 10mer - FRET | 10 | [Cy5]GGAACAAGGU | ACCUUGUUCC[Cy3] |
| ssRNA 10mer - Donor | 10 | ACCUUGUUCC[Cy3] | ACCUUGUUCC |
| ssRNA 10mer - Acceptor | 10 | [Cy5]GGAACAAGGU | GGAACAAGGU |
| ss RNA 10mer - FRET | 10 | [Cy5]GGAACAAGGU | [Cy3]GGAACAAGGU |

**Supplementary Table 6. Summary of correction factors and FRET measurements of ss and ds RNA in buffer and coacervate systems.**

| | System | | $E_{CT}$ (corrected FRET) |
|---|---|---|---|
| $\alpha$ (ss RNA 10mer) | In buffer | 0.073 ± 0.001 | -0.022 ± 0.001 |
| $\alpha$ (ss RNA 10mer) | ATP/$(Lys)_{10}$ | 0.068 ± 0.001 | -0.024 ± 0.003 |
| $\alpha$ (ss RNA 10mer) | $(Asp)_{30}/(Lys)_{30}$ | 0.088 ± 0.003 | -0.020 ± 0.017 |
| $\alpha$ (ds RNA 10mer) | In buffer | 0.085 ± 0.003 | 0.567 ± 0.015 |
| $\alpha$ (ds RNA 10mer) | ATP/$(Lys)_{10}$ | 0.066 ± 0.002 | 0.535 ± 0.076 |
| $\alpha$ (ds RNA 10mer) | $(Asp)_{10}/(Lys)_{10}$ | 0.070 ± 0.001 | 0.489 ± 0.077 |
| $\alpha$ (ds RNA 10mer) | $(Asp)_{30}/(Lys)_{30}$ | 0.108 ± 0.037 | 0.296 ± 0.047 |
| $\alpha$ (ds RNA 10mer) | $(Asp)_{100}/(Lys)_{100}$ | 0.072 ± 0.002 | 0.158 ± 0.031 |
| $\beta$ (ss RNA 10mer) | In buffer | 0.077± 0.001 | -0.022 ± 0.001 |
| $\beta$ (ss RNA 10mer) | ATP/$(Lys)_{10}$ | 0.044± 0.001 | -0.024 ± 0.003 |
| $\beta$ (ss RNA 10mer) | $(Asp)_{30}/(Lys)_{30}$ | 0.074 ± 0.007 | -0.020 ± 0.017 |
| $\beta$ (ds RNA 10mer) | In buffer | 0.067 ± 0.001 | 0.567 ± 0.015 |
| $\beta$ (ds RNA 10mer) | ATP/$(Lys)_{10}$ | 0.056 ± 0.001 | 0.535 ± 0.076 |
| $\beta$(ds RNA 10mer) | $(Asp)_{10}/(Lys)_{10}$ | 0.053 ± 0.003 | 0.489 ± 0.077 |
| $\beta$ (ds RNA 10mer) | $(Asp)_{30}/(Lys)_{30}$ | 0.054 ± 0.001 | 0.296 ± 0.047 |
| $\beta$(ds RNA 10mer) | $(Asp)_{100}/(Lys)_{100}$ | 0.096 ±0.003 | 0.158 ± 0.031 |

**Supplementary Table 7. Radiolabeled RNA Partitioning.** 5'-radiolabeled tRNAphe was added to coacervates, then centrifuged and the dilute phase was scintillation counted. % RNA in dilute phase calculated by dividing the average cpm/uL in the dilute phase by the average overall cpm/uL. Similar amounts of RNA were in all of the coacervate phases.

| Pair of Coacervates | Average cpm/µL in dilute phase | Standard Deviation (cpm/uL) | % RNA in dilute phase | % RNA in coacervate phase |
|---|---|---|---|---|
| $(Lys)_{10}$/ATP | 43.9 | 11.7 | 0.99 | 99.0 |
| $(Lys)_{10}/(Asp)_5$ | 319 | 52.6 | 4.16 | 95.8 |
| $(Lys)_{10}/(Asp)_{10}$ | 220. | 50.2 | 4.96 | 95.0 |
| $(Lys)_{30}/(Asp)_{30}$ | 238 | 84.15 | 5.37 | 94.6 |
| $(Lys)_{100}/(Asp)_{100}$ | 750. | 683 | 9.77 | 90.2 |

Values from samples that contained both coacervate and continuous phases were used to determine the fraction in each phase. Although the same amount of RNA was used in each sample, due to different amounts of radioactive decay these values were: 4430 cpm/µL for $(Lys)_{10}$/ATP, $(Lys)_{10}/(Asp)_{10}$, $(Lys)_{30}/(Asp)_{30}$, and 7680 cpm/µL for $(Lys)_{10}/(Asp)_5$, $(Lys)_{100}/(Asp)_{100}$
